# Site-Specific Knockdown of Microglia in the Locus Coeruleus Regulates Hypervigilant Responses to Social Stress in Female Rats

**DOI:** 10.1101/2022.10.03.509934

**Authors:** Brittany S. Pate, Samantha J. Bouknight, Evelynn N. Harrington, Sarah E. Mott, Lee M. Augenblick, Cora E. Smiley, Christopher G. Morgan, Brittney M. Calatayud, Gustavo A. Martinez-Muniz, Julian F. Thayer, Susan K. Wood

## Abstract

**Background:** Women are at increased risk for psychosocial stress-related anxiety disorders, yet mechanisms regulating this risk are unknown. Psychosocial stressors activate microglia, and the resulting neuroimmune responses that females exhibit heightened sensitivity to may serve as an etiological factor in their elevated risk. However, studies examining the role of microglia during stress in females are lacking.

**Methods:** Microglia were manipulated in the stress-sensitive locus coeruleus (LC) of female rats in the context of social stress in two ways. First, intra-LC lipopolysaccharide (LPS; 0 or 3μg/side, n=5-6/group), a potent TLR4 agonist and microglial activator, was administered. One hour later, rats were exposed to control or an aggressive social defeat encounter between two males (WS, 15-min). In a separate study, females were treated with intra-LC or intra-central amygdala mannosylated liposomes containing clodronate (m-CLD; 0 or 25μg/side, n=13-14/group), a compound toxic to microglia. WS-evoked burying, cardiovascular responses, and sucrose preference were measured. Brain and plasma cytokines were quantified, and cardiovascular telemetry assessed autonomic balance.

**Results:** Intra-LC LPS augmented the WS-induced burying response and increased plasma corticosterone and interleukin-1β (IL-1β). Further, the efficacy and selectivity of microinjected m-CLD was determined. In the context of WS, intra-LC m-CLD attenuated the hypervigilant burying response during WS as well as the accumulation of intra-LC IL-1β. Intra-central amygdala m-CLD had no effect on witness stress-evoked behavior.

**Conclusions:** These studies highlight an innovative method for depleting microglia in a brain region specific manner and indicate that microglia in the LC differentially regulate hypervigilant WS-evoked behavioral and autonomic responses.

**HIGHLIGHTS:**

- Intra-LC LPS augments behavioral and physiological responses to social stress
- Mannosylated liposomal clodronate site-specifically reduces microglial expression
- Microglia within the locus coeruleus regulate stress-evoked behavior in female rats

## 1. INTRODUCTION

Psychosocial disorders such as anxiety and depression have become increasingly prevalent, and notably, social conflict and isolation common to the COVID pandemic are considerable risk factors. These conditions lead to reduced quality of life, increased healthcare costs, and place significant burden on the affected individuals and society (Amos et al., 2018; Brody et al., 2018; Revicki et al., 2012). Social stressors, such as witnessing a traumatic event, often precipitate the onset of these debilitating disorders (Almeida, 2005; Feinstein et al., 2014; Kessler, 1997; Nicodimos et al., 2009; Post, 1992). Interestingly, females are more than twice as likely as males to suffer from these stress-related conditions (Asher et al., 2017; Brody et al., 2018; Hankin et al., 1998; Kessler et al., 1993). While the neural mechanisms underlying increased stress susceptibility among females are unclear, recent evidence highlights a critical role for stress-induced neuroimmune signaling as a causative factor (Bekhbat & Neigh, 2018; Finnell & Wood, 2018; Holmes et al., 2018; Moieni et al., 2015; Richards et al., 2018). In addition to alterations in neuroimmune signaling, social stress exposure also leads to major shifts in autonomic function marked by changes in cardiovascular function (Ring et al., 2002; Winters et al., 2000) to which women may be particularly susceptible (Vaccarino & Bremner, 2017). Furthermore, females report significantly greater depressed mood versus males when faced with a low-dose endotoxin challenge, despite mounting similar inflammatory responses. This indicates that females may be particularly susceptible to the psychological effects of inflammation (Moieni et al., 2015) and highlights a pathological mechanism that may underlie increased susceptibility to stress-related disorders in females. However, there is a significant lack of studies examining the regulatory role of microglia within discrete brain regions in response to social stress in females.

Inflammatory factors that accumulate in stress susceptible individuals (Brydon & Steptoe, 2005; Finnell & Wood, 2018; Maes et al., 1998; Wood et al., 2015) are capable of regulating neural activity. For example, pro-inflammatory cytokines increase neuronal activity within the locus coeruleus (LC), a brain nucleus that regulates the behavioral response of burying (Howard et al., 2008) that is classified as a hypervigilant behavior (Mikics et al., 2008), as well as autonomic responses to stressful stimuli (Wood & Valentino, 2017), key features associated with anxiety disorders (Grueschow et al., 2021; Morilak et al., 2005). In preclinical studies, pro-inflammatory cytokines are elevated in male mice that are susceptible to social stress (Hodes et al., 2014), and accumulate in the LC of stress-susceptible, but not resilient, male rats following social defeat stress (Finnell, Lombard, Melson, et al., 2017; Wood et al., 2015). Further, intra-LC microinjection of the pro-inflammatory cytokine interleukin (IL)-1β upregulates tonic and phasic activity of LC neurons (Borsody & Weiss, 2002a), and IL-6-dependent activation of LC neurons induces depressive-like behavior in rodents (Kurosawa et al., 2016). Importantly, certain patient subpopulations with depression and anxiety disorders also exhibit greater LC-norepinephrine (NE) activity that is associated with hypervigilance (Naegeli et al., 2018; Wong et al., 2000). Therefore, it is critical to understand if stress-induced neuroimmune signaling within the LC contributes to hypervigilant behaviors.

Neuroimmune responses become sensitized following repeated stress, largely through the key regulator high mobility group box-1 (HMGB-1), a pro-inflammatory molecule released by various cells including activated immune cells (Scaffidi et al., 2002; Wang et al., 1999). Importantly, in the context of psychosocial stress, microglia release HMGB-1 to promote this stress-induced sensitization (Frank et al., 2018; Frank et al., 2015; Weber et al., 2015; H. Zhang et al., 2019). Therefore, microglia may serve as a therapeutic target for the treatment of stress-related disorders (Calcia et al., 2016; Jia et al., 2021; Mondelli et al., 2017; Weber et al., 2019). Several preclinical studies in males have demonstrated that inhibiting macrophages via minocycline or depleting microglia using colony stimulating factor 1 receptor (CSF1R) antagonist, alleviates stress-induced inflammation and ultimately improves depressive- and anxiety-like behaviors (Kobayashi et al., 2013; Wang et al., 2018; C. Zhang, A. V. Kalueff, et al., 2019; C. Zhang, Y. P. Zhang, et al., 2019). These studies identify a critical role for immune cells in stress-induced behavioral maladaptation in males; however, these treatments are administered via diet, resulting in global changes in macrophages and microglia. To date, no studies have utilized microglial depletion within discrete brain regions *in vivo* to determine how microglia in stress-sensitive brain regions facilitate physiological and behavioral responses to social stressors. The recent development of mannosylated liposomal clodronate (m-CLD) may provide the opportunity to target microglia within specific brain regions. Due to mannose-targeting, the mannosylated liposomes are taken up specifically by microglia via phagocytosis, facilitating the uptake of clodronate into the cell. Once in the cytosol, clodronate disrupts mitochondrial respiration, ultimately leading to cell death (Lehenkari et al., 2002; Torres et al., 2016; Umezawa & Eto, 1988).

Given the increased risk of stress-related disorders in females, the current studies investigated the role of intra-LC microglia in behavioral and autonomic responses to social stress exposure in female rats. While a lack of ethologically relevant models of social stress in female rats previously limited our mechanistic understanding of social stress susceptibility among females, our lab and others have demonstrated that vicarious/witness stress (WS), a modified version of the resident-intruder model, is an effective social stress paradigm in both male (Finnell, Lombard, Padi, et al., 2017; Patki et al., 2014; Sial et al., 2016; Warren et al., 2020) and female rodents (Finnell et al., 2018; Iniguez et al., 2018; Warren et al., 2020). We determined that WS promotes lasting increases in central IL-1β, hypervigilant burying, reduced sucrose preference, immobility during the forced swim test, and autonomic deficits in female rats (Finnell et al., 2018). In addition to prior work from our lab identifying the neuroimmune response to WS, other variations of WS have demonstrated a neuroinflammatory response (Ataka et al., 2013; Lukkes et al., 2017). The current study expanded on these findings to determine 1) if the toll like receptor-4 (TLR4) agonist lipopolysaccharide (LPS)-induced activation of microglia selectively within the LC prior to an acute social stress exposure resulted in an exaggerated hypervigilant behavioral response, 2) if m-CLD-induced reduction of microglial expression, injected into the LC prior to repeated WS would attenuate the expression of hypervigilant behaviors and reduced sucrose preference, and 3) if partial microglial depletion in the LC also regulated stress-induced hypervigilant autonomic responses. Further, a separate subset of rats was subjected to m-CLD treatment in the central nucleus of the amygdala (CeA) to determine the specificity of effects observed compared with LC microglial knockdown.

## 2. METHODS AND MATERIALS

### 2.1 Animals

Female Sprague Dawley rats (8-9 weeks of age upon arrival, experimental rats; Charles River, Raleigh, NC), male Sprague Dawley rats (275-300g, intruders; Charles River), and male Long-Evans retired breeders (650-850g, residents; Charles River or Envigo, Indianapolis, IN) were individually housed in standard cages within a climate-controlled room on a 12-hour light/dark cycle. Rats were given *ad libitum* access to food and water. These studies were approved by the University of South Carolina’s Institutional Animal Care and Use Committee and all experiments followed the National Research Council’s *Guide for the Care and Use of Laboratory Animals*.

### 2.2 Study Design

Four separate studies were conducted. Study A was designed to understand the role of LPS-induced TLR4 microglial activation within the LC on behavioral and physiological responses to WS (**Figure 1A**). Further, a series of studies were conducted to characterize the efficacy and selectivity of m-CLD in depleting LC microglia (Study B; **Figure 2A**) and the subsequent impact on behavioral and autonomic responses to WS (Study C; **Figure 3A**). Finally, in a design comparable to study C, m-CLD was infused into the CeA (Study D; **Figure 4A**).

**Figure 1.**
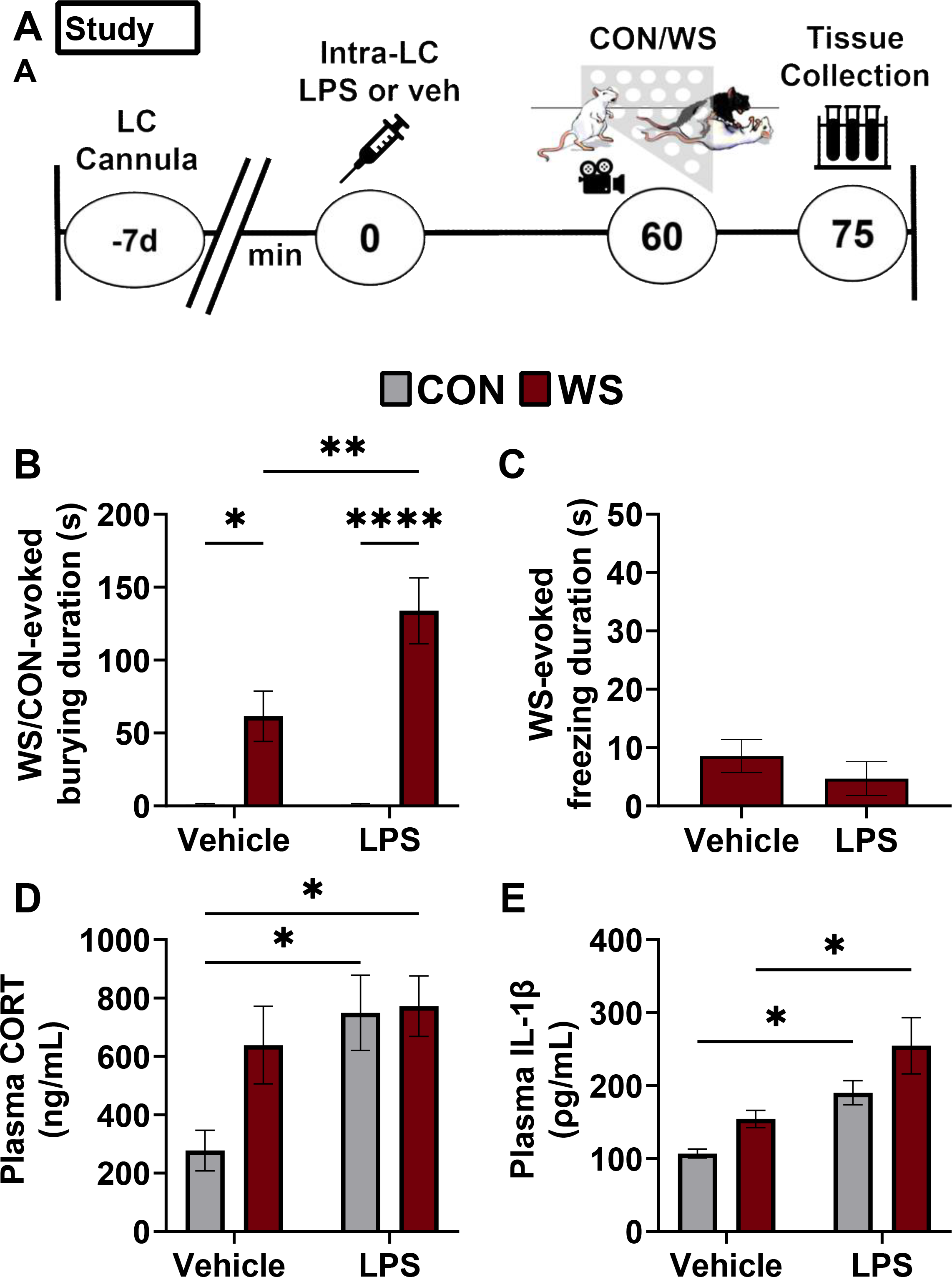
Effects of intra-locus coeruleus (LC) lipopolysaccharide (LPS) on control or witness stress (WS)-evoked behavior and plasma Cort and IL-1β responses. **(A)** This panel depicts the intra-LC LPS study design (**Study A**). Seven days (d) following surgical implantation of an indwelling bilateral cannula into the LC, rats were briefly anesthetized (isoflurane time from anesthesia induction to awake ranged from 40-107 sec) and microinjected with saline vehicle or lipopolysaccharide (LPS, 3μg/side) into the LC 60 min prior to an acute exposure to control handling (CON) or a 15-min period of witness stress (WS) immediately prior to tissue collection (CON + vehicle (*n* = 5); WS + vehicle (*n* = 5); CON + LPS (*n* = 6); WS + LPS (*n* = 8)). **(B)** Rats spent significantly more time burying during WS compared to their respective controls, regardless of treatment (\**p* < 0.05, CON + vehicle (n=5) vs WS + vehicle (n=5); \*\*\*\**p* <0.0001, CON + LPS (n=6) vs WS + LPS (n=5)), and WS-induced burying is further increased by LPS treatment (\*\**p* < 0.01, WS + vehicle vs WS + LPS). **(C)** WS-induced freezing behavior is not affected by LPS treatment (unpaired *t*-test, *t_8_* = 0.96 *p* = 0.37). **(D)** Plasma corticosterone (CORT) levels were increased by LPS and WS to a similar extent by intra-LC LPS and WS (main effect of stress *F_1,15_* = 6.7, *p* = 0.12; significant effect of drug *F_1,15_* = 6.67, *p* = 0.02). Independent of stress, CORT was also increased by intra-LC LPS treatment in CON rats (\**p* < 0.05 CON + vehicle (n=5) vs CON + LPS (n=6)). However, intra-LC LPS combined with WS did not produce a greater CORT response than WS (*p* = 0.88, WS + vehicle (n=4) vs WS + LPS (n=5)) or intra-LC LPS (*p* = 0.99, CON + LPS (n=6) vs WS + LPS (n=5)) alone. **(E)** WS and intra-LC LPS produced an exaggerated plasma IL-1β response (effects of stress: F_1,18_ = 7.2, *p* = 0.02; and drug: F_1,18_ = 19.4, *p* = 0.0003). In the absence of LPS, WS exposure elicited a modest, yet non-significant increase in plasma IL-1β (*p* = 0.40; CON + vehicle (n=5) vs. WS + vehicle (n=6)) likely due to plasma collection occurring immediately following the 15-min WS exposure. However, intra-LC LPS in the absence of WS produced an increase in plasma IL-1β (\**p* = 0.04 CON + vehicle (n=6) vs CON + LPS (n=5)) while WS combined with intra-LC LPS treatment produces a greater peripheral IL-1β response than WS alone (\**p* = 0.02 WS + vehicle (n=6) vs WS + LPS (n=5).

**Figure 2.**
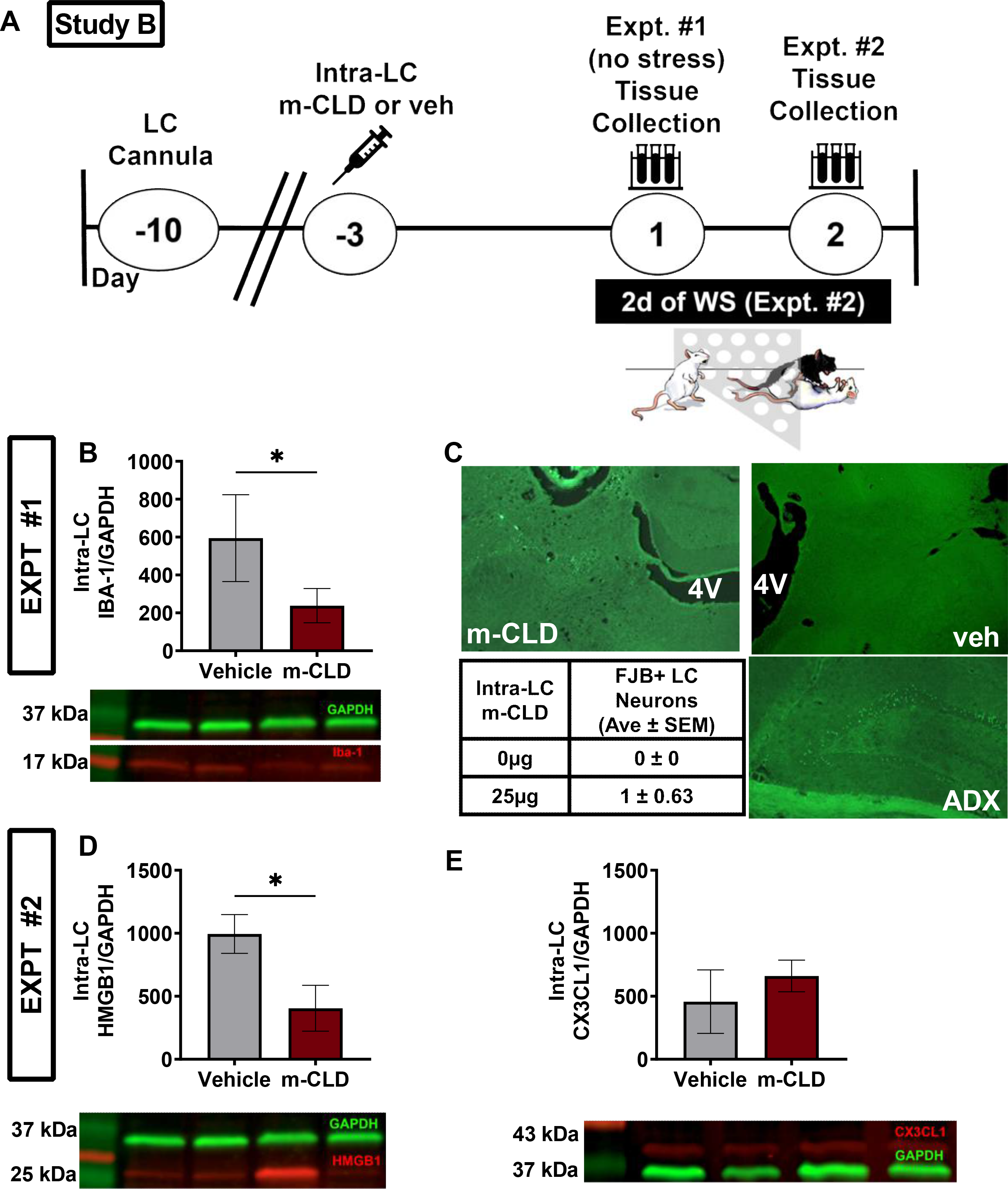
Characterization of the effect of discrete injections of mannoslyated clodronate (m-CLD) into the LC. **(A)** Timeline in days of the two experiments conducted to assess the efficacy, selectivity, and potential compensatory effects of intra-LC m-CLD (25μg/side). Experiment 1 collected tissue under resting conditions from vehicle or m-CLD treated rats 3 days following the injection into the LC to confirm **(B)** effective knockdown of microglia within the LC (IBA-1 positive cells; *t_5_=2.9, *p*<0.05). Fluorojade B (FJB), a stain used to detect degenerating neurons was used to determine if m-CLD (25μg) elicited neuronal damage **(C)**. A representative fluorescent photomicrograph of tissue collected 3 days following bilateral infusion of intra-LC m-CLD (25μg; top left) or vehicle (0μg; top right). The 4^th^ ventricle (4v) is marked in these images. Data from vehicle-treated m-CLD infused rats were processed using FJB and quantified in the bottom left table (*t_6_=0.87, p* =0.42). Hippocampal tissue from an adrenalectomized (ADX) rat was used as a positive control (bottom right) as this is known to induce cell death within the hippocampus. Because it is known that FJB also reacts with activated microglia (Damjanac et al., 2007), tissue was double labeled for IBA-1 and FJB and only FJB-positive / IBA-1-negative cells were considered markers of neuronal degeneration. A representative cluster of FJB-positive cells, IBA-1-positive cells, and the merged image is shown in **Figure S5**. Next, two major factors, HMGB1 and CX3CL1, could function to elicit a compensatory effect to overcome the deficit induced by fewer numbers of microglia. Therefore, in experiment #2, tissue from vehicle or m-CLD treated rats was collected following the 2^nd^ exposure to WS. **(D)** m-CLD-induced decreases in microglial expression resulted in the expected reduction in HMGB1 expression in WS-exposed rats compared with vehicle treated WS rats (HMGB1/GAPDH; *t_11_* =2.49 \**p* = 0.03), proving that an HMGB1-related compensatory response was not occurring in existing microglia. (**E**) Further, it was established that neighboring cells did not withdraw their inhibitory control over microglia via CX3CL1 to compensate for reduced microglial tone (CX3CL1/GAPDH; *t*_11_=1.79 *p* = 0.10). Cropped images from representative western blots are shown to illustrate expression of relevant proteins below their respective graphs. Full blot images are provided in the supplemental (**Figure S7**). All data are provided as mean ± SEM and unpaired *t* tests were used for all analyses.

**Figure 3.**
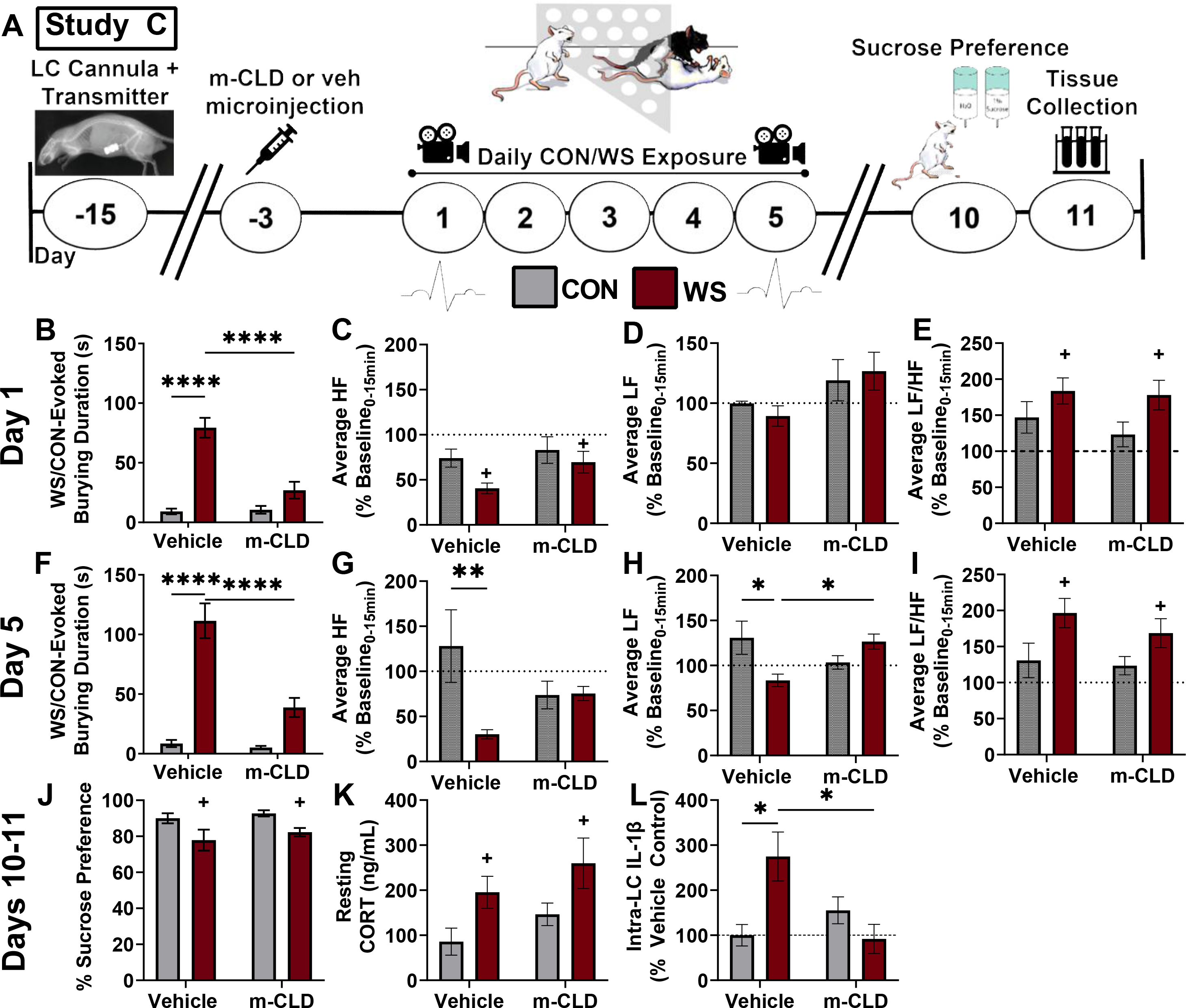
Effects of intra-LC m-CLD on witness stress (WS)-induced behavior and heart rate variability (HRV) during WS exposure. **(A)** This panel depicts the study timeline for the intra-LC m-CLD study which was used to determine the impact of reducing the microglial expression within the LC on the consequences of repeated WS exposure. The behavioral response to WS was striking among vehicle treated rats, with robust expression of burying during WS on day 1 (**B**; \*\*\*\**p* < 0.0001 CON + vehicle (n=13) vs. WS + vehicle (n=13)) and day 5 (**F**; \*\*\*\**p* < 0.0001 CON + vehicle (n=13) vs. WS + vehicle (n=13)). Importantly, intra-LC m-CLD blunted the WS-induced increase in WS-evoked burying on day 1 (**B**; *\*\*\*\*p* < 0.0001 WS + vehicle vs. WS + m-CLD (n=10)) and day 5 (**F**; \*\*\*\**p* < 0.0001 WS + vehicle (n=13) vs. WS + m-CLD (n=10)). Spectral analysis of HRV was used to determine the effects of WS and intra-LC m-CLD on autonomic function. WS resulted in an upward shift in the low frequency(LF) to high frequency (HF) ratio (LF/HF) on day 1 **(C**; *F*_1,22_ = 4.77, *p* 0.04), and following repeated WS exposure on day 5 (**G**; effect of stress *F_1,23_* = 6.41, *p* =0.02) with no significant post hoc analyses. Interestingly, LF, indicative of sympathetic drive was increased slightly by m-CLD compared to vehicle with no significant post-hoc analyses on Day 1 (**D**; effect of drug: *F_1,23_* = 5.00, *p* = 0.03), and a stress x drug interaction emerged following repeated stress on day 5 (**H**; F1,23 = 11.72, p = 0.002). Further, HF, primarily indicative of parasympathetic drive, or vagal tone, was reduced by WS compared to control on day 1 (**E**; effect of stress: *F_1,21_* = 5.17, *p* = 0.03) with no effect of m-CLD. However, following repeated stress, m-CLD blocks this vagal withdrawal induced by WS (**I**; effect of stress: *F_1,22_* = 7.49, *p* = 0.01; stress x drug interaction: *F_1,22_* = 7.99, *p* = 0.01). Sucrose preference was modestly reduced among WS rats compared with controls (**J;** effect of stress *F_1,43_* = 8.05, \**p* = 0.01). However, this stress-induced reduction in sucrose preference was not altered by intra-LC m-CLD treatment. Similarly, resting plasma CORT levels are increased in rats with a history of repeated WS exposure (**K**; CON + vehicle (n=5); CON + m-CLD (n=4); WS + vehicle (n=6); WS + m-CLD (n=5); effect of stress *F_1,16_* = 7.73 \**p* = 0.01), and was not affected by intra-LC m-CLD treatment. Finally, these studies confirmed that IL-1β expression within the LC is persistently increased in rats with a history of WS as evidenced six days after the cessation of stress (**L**;*\*p*< 0.05 CON + vehicle (n=5) vs. WS + vehicle (n=6)). Importantly, this effect was attenuated by intra-LC m-CLD (L \**p* < 0.05 WS + m-CLD (n=5) vs WS + vehicle (n=5)). Significant main effects of stress are indicated by “+” while significant post hoc analyses are denoted on figure (*p < 0.05, **p<0.01).

**Figure 4.**
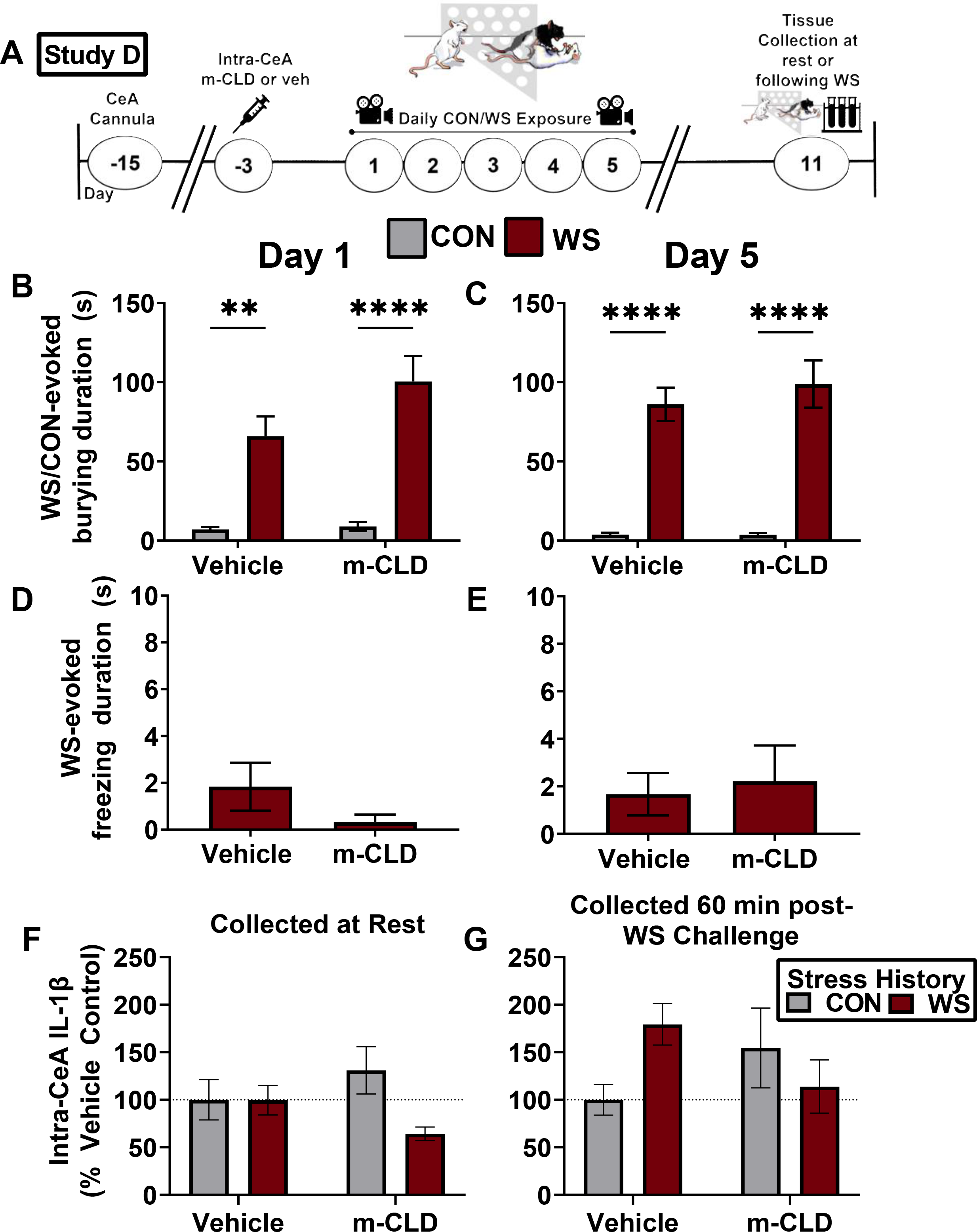
Effects of Intra-CeA m-CLD on microglial expression and WS-induced accumulation of brain IL-1β. (**A**) Timeline for Study D examining the effect of intra-CeA m-CLD on WS-induced neuroimmune and behavioral responses (**Study D**). Study D was conducted identically to Study C with the exception of the cardiovascular transmitter surgery and sucrose preference testing. **(B)** WS induced robust burying behavior on day 1 (\*\**p* <0.005 CON + vehicle (n=13) vs. WS + vehicle (n=12); \*\*\*\**p*<0.0001 CON + m-CLD vs. WS (n=13) + m-CLD (n=13) and **(C)** day 5 (*\*\*\*\*p* < 0.0001 CON + vehicle (n=13) vs. WS + vehicle (n=12); \*\*\*\**p*<0.0001 CON + vehicle (n=13) vs. CON + m-CLD (n=13)). Among WS rats, m-CLD did not regulate freezing duration on Day 1 **(D**; unpaired *t* test, *t_24_* = 1.41, *p* = 0.17)or Day 5 **(E). (F)** There was no significant effect of WS or m-CLD treatment on resting levels of IL-1β expression six days after the last WS/con exposure. **(G)** However, in a subset of WS and CON rats that were re-exposed to WS 60 min prior to sacrifice, a history of WS resulted in a sensitized WS-induced IL-1β response in vehicle-treated witnesses, but not m-CLD treated witnesses (stress x drug interaction: *F*_1,22_ = 5.31, *p* = 0.031).

### 2.3 Surgery

In Studies A, B and C, female witnesses and controls were implanted with indwelling bilateral cannulas (26ga., P1 Technologies, Roanoke, VA) into the LC (caudal: −3.9 mm from bregma; lateral: ± 1.1 mm; and dorsal-ventral: −5.1 mm from skull, nose at a 30° angle). In Study C, a subset of witnesses and controls were also implanted with HD-S11 radiotelemetric transmitters (Data Sciences International, St. Paul, MN) in the modified lead II configuration as previously published (Finnell, Lombard, Padi, et al., 2017; Wood et al., 2012). In Study D, female witnesses and controls were implanted with indwelling bilateral cannulas aimed at the CeA (caudal: −2.3 mm from bregma; lateral: ± 4.1 mm; and dorsal-ventral: −6.5 mm from skull). All rats were given standard post-operative care (Flunazine, 0.25 mg/kg) and allowed at least 7-10 days of recovery prior to experimental manipulations.

### 2.4 Drug Information

#### Study A – LPS

LPS was selected as an agent to induce microglial activation because LPS increases microglial activity through TLR4 in a similar fashion to HMGB1. Importantly, preliminary data suggests that our WS model produces prolonged increases in HMGB1 in the LC as measured 6 days post stress/control (unpaired *t* test, *p* = 0.06; **Figure S1**), highlighting the relevance of selecting LPS. LPS (Sigma-Aldrich, St. Louis, MO) was dissolved in 0.9% sterile saline at a concentration of 3μg/μL. Witnesses and controls received 1μL of either 0μg/μL or 3μg/μL LPS microinjected bilaterally into the LC. This dose was based on pilot studies demonstrating 3μg/μL would induce stress-evoked behaviors and elicit an inflammatory response without producing sickness behaviors (**Figure S2**).

#### Studies B, C and D – Mannosylated Liposomal Clodronate (m-CLD)

Rats were administered mannosylated liposomes containing clodronate (m-Clodrosome® (m-CLD); Encapsula NanoSciences, Brentwood, TN) or empty mannosylated liposomal vehicle (veh; m-Encapsome®; Encapsula Nanosciences). Each female rat was microinjected with m-CLD (25μg/side) or vehicle (0μg/side) directly into the LC (studies B, C;) or the CeA (study D). This dose of m-CLD was chosen based on pilot studies reported in this manuscript demonstrating m-CLD sufficiently reduced the local microglial population by ~50% without causing neuronal damage. Additional studies confirmed that the injection did not spread beyond our targeted region using a mannosylated fluorescent liposome (Encapsula Nanosciences, **Figure S4**).

### 2.5 Witness Stress (WS)

Female witnesses were placed behind a clear, perforated, acrylic partition in a protected region of a novel resident’s cage where they observed the auditory, visual, and olfactory cues of social defeat between the resident and an intruder for 15 minutes as described in detail in Finnell et al., 2018 (Finnell et al., 2018). Controls were briefly handled and placed back into their home cage. This method of control handling was used because it was previously confirmed that handling induced the same response as being placed behind a partition in a male Sprague-Dawley’s cage ((Finnell et al., 2018) see supplement).

### 2.6 Behavioral Measurements

#### 2.6.1 Behavior During WS and Control

WS and control sessions were video-recorded and behaviors including rearing, freezing, and stress-evoked burying were manually quantified retrospectively by a blinded experimenter using ANY-maze (Stoelting Co., Wood Dale, IL) and validated by a second blinded scorer. Behaviors were measured during exposure to social defeat for WS rats and immediately following handing for control rats and were defined as the following: stress-evoked burying - frantic, non-exploratory digging or pushing of the bedding; freezing - complete immobility except for respiratory movements; and rearing - any time at which the rat’s front paws were simultaneously raised from the bedding with or without the hindlimbs extended, in the absence of grooming, eating, or drinking. Analyses included the latency to the first occurrence of each behavior as well as the total duration of each behavior. Freezing for controls is not reported due to a high level of sleeping during the recording period that confounded freezing measures.

#### 2.6.2 Assessment of Sickness Behavior

In Study A, to confirm that the intra-LC LPS dose range used in this study did not elicit sickness, several sickness behaviors that have been shown to be regulated by pro-inflammatory cytokines were quantified for the 0-3μg doses of LPS that were used in the study (Bassi et al., 2012; Chaskiel et al., 2021; Dantzer, 2001; Foster et al., 2021; Grossberg et al., 2011; Harden et al., 2011). During resting conditions, following control handling, the following behaviors were evaluated as an indication of sickness: ragged fur, absence of cage exploration, squinted eyes not associated with sleep, and lethargy (general withdrawal and lack of interest in the environment). During witness stress, rats were assessed for the following behaviors as an indication of sickness: absent grooming, non-reactive to resident-intruder aggression, sleeping, and hunched/curled posture. For each sickness behavior displayed at any time during the testing period, one point was tallied. To be included in the study, rats had to have fewer than three sickness points. Our dose response curve was originally designed to test 0-10μg, however three rats were treated with the high dose and displayed robust sickness behaviors, so it was determined that our highest dose should be limited to 3μg. Importantly, no rats in the 0, 1, and 3μg groups were considered sick, and no exclusions had to be made based on these criteria (see **Figure S2E**).

#### 2.6.3 Sucrose Preference Test

To assess the impact of WS and m-CLD treatment on anhedonia, a two-bottle-choice sucrose preference test was performed in Study C. The sucrose preference test was conducted over a 72-hour period using three phases with food freely available during all phases as previously published (Finnell, Lombard, Melson, et al., 2017; Finnell et al., 2018; Mao et al., 2014; Wood et al., 2015). Phase 1: rats were acclimated to two water bottles in their home cage for the first 24 hours. Phase 2: For the next 24 hours, rats had free access to two standard bottles filled with 1% sucrose. Phase 3 (test day): Rats were water deprived for 10 hours until the start of the dark cycle (9am – 7pm), when two bottles, one with water and one with 1% sucrose, were placed in the cage for 3 hours (7pm-10pm). Total weight of liquid consumed from each bottle was calculated and preference was determined by dividing the weight of 1% sucrose consumed by all liquid consumed. The location (right vs left) of the sucrose bottle was counterbalanced between rats and the order of the bottles was switched at 8pm. Rats were excluded if they exhibited sucrose aversion indicated by preference below 50% in the pre- and post-stress sucrose preference test (exclusions: WS + m-CLD n=1; WS + vehicle n=1; CON + m-CLD n=0; CON + vehicle n=0).

### 2.7 Cardiovascular Telemetry

In Study C, radiotelemetric implants allowed for cardiovascular data acquisition during WS. Cardiovascular data were acquired using Ponemah Acquisition software (Data Sciences International, St. Paul, MN) and analyzed via the Ponemah physiology platform (Data Sciences International) as previously published (Finnell, Lombard, Padi, et al., 2017).

#### Heart Rate Variability (HRV) Analyses

Frequency domain analysis of heart rate variability (HRV) was conducted to determine the spectral power within the low-frequency domain (LF), a nonspecific index of sympathetic activity, and the high-frequency domain (HF) an index of parasympathetic (vagal) activity (Berntson et al., 1997). The LF/HF ratio, an estimate of sympathovagal balance (Malliani et al., 1991), was used to assess autonomic responses on days 1 and 5 of WS/control. Continuous HRV data were collected during the light phase on day 1 and day 5 of WS/control. Light cycle baseline values were recorded while rats remained undisturbed in their home cages prior to the start of WS/control manipulations. Continuous measurements during the 15 minutes of WS or immediately following control handling were obtained and used to assess WS- or control handling-induced changes in HRV. The LF/HF ratio during stress was calculated as a percent of baseline and data are presented as the average percent baseline during WS/CON exposure(Graph Pad Prism 9.1.1). Telemetry data also provided measures of heart rate (HR) and mean arterial blood pressure (MAP). Additional CV measures of cardiac output (CO), total peripheral resistance (TPR), and stroke volume were derived from these telemetry data as described in the supplemental methods.

### 2.8 Tissue Collection and Processing

Rats were briefly anesthetized with isoflurane prior to decapitation. Brains were immediately dissected, flash-frozen in 2-methylbutane, and stored at −80°C. Brains were sliced coronally and bilateral punches of the LC were taken using a 1mm diameter tissue biopsy punch (Harvard Apparatus, Holliston, MA). Histological assessment verified accuracy of punch placement and cannula location (**Figure S3**). Punches were homogenized and assessed for protein concentration using a Pierce Bicinchoninic Acid (BCA) assay (Thermo Scientific, Rockford, IL) per manufacturer’s protocol. Briefly, each sample was mixed with 100 μL of zirconium oxide beads and 200 μL of lysis buffer consisting of 137mM NaCl, 20 mM Tris, 1% Ipegal, 10% glycerol, and 1x protease inhibitor cocktail. Samples were placed into a Bullet Blender for 3 minutes on speed 8 at 4°C followed by centrifugation at 1400 rcf for 15 minutes at 4°C. Resulting homogenate for each sample was mixed in a 1:4 dilution with phosphate buffered sodium azide (PB-azide). Standards and samples (25 μL) were loaded into a 96-well plate. 200 μL of working reagent was added to each well and the plate was incubated at 37°C for 30 minutes before reading at 562 nm. homogenates containing 20 μg of protein were aliquoted and stored at −80°C. Trunk blood was collected in EDTA-lined tubes (BD Vacutainer, Franklin Lakes, NJ), centrifuged in 4°C at 1600 rcf for 15 minutes to extract plasma, and stored at −80°C.

#### 2.8.1 Histology

Immediately prior to and following punch acquisition, a series of 30μm thick slices were obtained and mounted onto slides for histological verification of punch placement. Slides were stained using Neutral Red (1.2g in 0.1M acetic acid), dehydrated in a series of ethanol baths, and fixed in histoclear (National Diagnostics, Atlanta, GA). Rats with both cannulas outside the brain region of interest were excluded from the study and punches found to be more than 30% outside the region of interest were also excluded from the study.

### 2.9 Molecular Analyses

#### 2.9.1 Corticosterone (CORT) and Cytokine Analyses

Plasma acquired from Studies A, C, and D were analyzed for CORT using a commercially available CORT ELISA kit (Enzo Life Sciences, Farmingdale, NY) performed per manufacturer’s protocol. In Study A, plasma levels of IL-1β were measured using a commercially available ELISA kit (R&D Systems, Biotechne, Minneapolis, MN) while brain cytokine analyses for Studies C and D were conducted using a custom 10-plex Bioplex (Bio-rad Laboratories, Hercules, CA) as previously described (Finnell, Lombard, Melson, et al., 2017) and previously validated in females (Finnell et al., 2018).

#### 2.9.2 Western Blotting

Following homogenization and BCA analysis, 20μg of LC homogenate (Study B) was subjected to western blot analysis. Gel electrophoresis was run using 4-20% gradient Mini-PROTEAN gels (Bio-Rad Laboratories, Hercules, CA) with a Chameleon Duo molecular weight ladder (LiCOR, Lincoln, NE). Protein was then transferred onto Infrared PVDF membranes (Millipore, Burlington, MA) and membranes were blocked in Immobilon Blok Solution (Millipore, diluted at 1:1 in 0.1M PBS) for 1 hour. PVDF membranes incubated overnight at 4°C with the following primary antibodies diluted in 1:1 Blok/0.1M PBS + 0.2% Tween-20 : mouse anti-GAPDH (1:1000, Santa Cruz, Santa Cruz, CA), rabbit anti-CX3CL1 (1:1000, Abcam), rabbit anti-HMGB1 (1:5000, Abcam), and goat anti-Iba1 (1:3000 Abcam). After washing, membranes incubated at room temperature for 2 hours with secondary antibodies (LiCor): IRDye 800CW donkey anti-mouse, IRDye 680LT donkey anti-rabbit (1:15,000), and IRDye 680LT donkey anti-goat (1:15,000) diluted in in 1:1 Blok/0.1M PBS + 0.2% Tween-20+ 0.01% SDS. Membranes were scanned using the LiCor Odyssey CLX System and quantified using Image Studios software (LiCor). Protein bands for each sample were normalized to their respective GAPDH band.

### 2.10 Statistical Analysis

Statistical outliers were identified as values greater than two standard deviations from the mean. Analyses were conducted using GraphPad Prism 9 software (GraphPad Software, La Jolla, CA). Data were subjected to analysis via 2-way ANOVA (α = 0.05) to determine effects of stress and drug. All ANOVAs were followed by Tukey’s post hoc analysis. Summary statistics for all 2-way ANOVA are presented in supplemental tables. Data from m-CLD characterization studies were subjected to unpaired *t* tests. Statistical statements/symbols in the figures are explained in the corresponding figure legends.

## 3. RESULTS

### 3.1 Intra-LC LPS Augments WS-Induced Anxiety-like Behavior (Study A)

Vehicle-treated witnesses (WS + vehicle; n=5) exhibited a significantly greater burying duration compared to controls (CON + vehicle (n=5); *p* = 0.03 vs. WS + vehicle), and WS-evoked burying behavior was further enhanced by intra-LC LPS selectively in the WS-exposed rats (WS + LPS (n=5) vs. CON + LPS (n=6), *p* < 0.0001). (**Figure 1B**; effect of stress *F_1,17_* = 7.19, *p* = 0.02; drug *F_1,17_* = 51.53, *p* < 0.0001; and interaction *F_1,17_* = 7.20, *p* = 0.02). Freezing, was evaluated in the witnesses, and was unaffected by intra-LC LPS (**Figure 1C**; WS + vehicle (n=5) vs. WS + LPS (n=5), unpaired *t*-test *t*_*15*._ = 0.83, *p* = 0.42). Importantly, these effects were not driven by differences in the severity of stress exposure as the average number of attacks towards the intruder did not differ between groups (unpaired *t*-test *t*19 = 1.07, *p* = 0.30; **Supplemental Figure S10A**). Rearing behavior occurs equally during control and WS exposures (**Figure S11E**; CON + vehicle (n=5) vs. WS + vehicle (n=5), *p* = 0.7), but among controls, LPS appears to reduce rearing behavior during control (**Figure S11E**, CON + vehicle (n=5) vs CON + LPS (n=5), *p* = 0004).

### 3.2 Effects of intra-LC LPS on WS-Induced Changes in CORT and IL-1β (Study A)

Intra-LC LPS increases circulating CORT levels regardless of stress condition (**Figure 1D;** effect of drug *F_1,15_* = 6.67, *p* = 0.02). Additionally, there was a clear trend towards elevated circulating CORT in response to WS exposure (effect of stress *F_1,15_* = 2.68, *p* = 0.12). However, prior treatment with intra-LC LPS did not further enhance the WS-evoked peripheral CORT response (**Figure 1D**; WS + vehicle (n=4) vs. WS + LPS (n=5) *p* = 0.88). In contrast, intra-LC LPS did augment the WS-induced increase in plasma IL-1β (effect of stress *F*_1,18_=7.68, *p* = 0.013; effect of drug *F*_1,18_ = 18.86, *p* = 0.0004), with the combination of WS + LPS producing the greatest effect (**Figure 1E;** WS + vehicle (n=6) vs WS + LPS (n=5) *p* = 0.02).

### 3.3 Characterization of Intra-LC m-CLD (Study B)

Two experiments were performed to characterize the effect of intra-LC m-CLD (25μg/side) (**Figure 2A**). Intra-LC m-CLD reduced LC microglial expression as shown by reduced Iba-1 expression within the LC 3 days following m-CLD treatment (**Figure 2B**; unpaired *t* test, *t_5_* = 2.9, \**p* = 0.03 vehicle (n=4) vs m-CLD (n=4)). This effect was further confirmed by immunohistochemistry in brains collected at the same timepoint (Supplemental **Figure S5**). We also demonstrated that this dose of m-CLD injected bilaterally into the LC did not induce neuronal damage as indicated by a lack of Fluorojade B (FJB) positive cells (**Figure 2C**). It should be noted that studies have identified that while FJB is a well-recognized fluorescent marker used for histological staining of degenerating neurons (Varvel et al., 2016), it has also been shown to mark activated microglia (Damjanac et al., 2007). Therefore, slices were stained for both FJB and IBA-1 and only FJB positive/IBA-1 negative cells were counted as dying neurons (**Figure S6**).

To determine if compensatory molecular effects occurred in CLD-treated rats in the face of WS, we assessed HMGB-1, a factor known to sensitize microglia, and Cx3CL1, an immunomodulatory chemokine with the potential to exert inhibitory effects on microglia. In a subset of rats that were treated with intra-LC m-CLD and underwent two consecutive days of WS, HMGB-1 remained reduced compared with vehicle treatment (**Figure 2D**, unpaired *t* test, *t_11_* = 2.5, \**p* = 0.03 vehicle (n=7) vs. m-CLD (n=8)) as expected from the reduced microglial levels. Importantly, no changes occurred in Cx3CL1 compared to vehicle treatment (**Figure 2E**, unpaired *t* test, *t_11_* = 1.8, *p* = 0.1 vehicle (n=7) vs m-CLD (n=8)). Together, these findings provide support for the assumption that selective depletion of microglia in the LC are responsible for any effects observed from m-CLD administration in these studies.

### 3.4 Intra-LC m-CLD Regulates WS-Evoked Burying (Study C)

In Study C (**Figure 3A**), anxiety-like burying duration was increased during WS on the first exposure (day 1, **Figure 3B**, effect of stress: *F_1,39_* = 55.56, *p* < 0.0001; CON + vehicle, n=13; CON + m-CLD, n=13; WS + vehicle, n=13; WS + m-CLD, n=10) and following repeated exposure (day 5, **Figure 3F**, effect of stress: *F_1,40_* = 59.67, *p* < 0.0001; CON + vehicle, n=13; CON + m-CLD, n=13; WS + vehicle, n=13; WS + m-CLD, n=10), and this effect was blunted by intra-LC m-CLD on both the first and fifth exposure (Day 1: effect of drug: *F_1,39_* = 19.26, *p* < 0.0001; Day 5: effect of drug: *F_1,40_* = 18.54, *p* < 0.0001). In fact, the effect of m-CLD was so effective that WS did not significantly increase burying over m-CLD treated controls (stress-evoked burying day 1: *p* = 0.26, CON + m-CLD vs. WS + m-CLD and stress-evoked burying day 5: *p* = 0.08, WS + vehicle vs. WS + m-CLD). Importantly, these marked group differences cannot be attributed to differences in the number of attacks the witnesses in each group observed as this was consistent across treatment groups (unpaired *t* test: *t_8_* = 0.15, *p* = 0.88; **Figure S10B**). Further, m-CLD induced reduction of intra-LC microglia did not impact freezing behaviors during WS on Day 1 WS + vehicle (1.9 s ± 1.1) vs. WS + m-CLD (0.49s ± 0.49); unpaired *t*-test, *t_18_* = 0.83, *p* = 0.42) or Day 5 (WS + vehicle (0.9 s ± 0.67) vs. WS + m-CLD (2.4 s ± 1.89); unpaired *t*-test *t_18_* = 0.91, *p* = 0.38). Rearing behavior was not affected by stress nor m-CLD on Day 1 (**Figure S11A**), but during day 5, rearing was reduced among controls compared to witnesses treated with m-CLD (**Figure S11B**; CON + m-CLD vs. WS + m-CLD, *p* = 0.0008).

### 3.5 Intra-LC Microglia Facilitate Vagal Withdrawal Following Repeated WS Exposure

Because the LC projects inhibitory input to the dorsal motor nucleus of the vagus, spectral analyses were used to determine the effects of WS and m-CLD on vagal tone (HF). WS results in vagal withdrawal as indicated by reduced HF during day 1 of WS (**Figure 3C**; effect of stress: *F_1,21_* = 5.17, *p* = 0.03) with no effect of m-CLD. However, following repeated WS exposure, m-CLD blocked this WS-evoked vagal withdrawal (**Figure 3G**; stress x drug interaction: *F_1,22_* = 7.99, *p* = 0.01). However, a compensatory increase in LF was observed following repeated WS on day 5 (**Figure 3H**; *F_1,23_* = 11.72, *p* = 0.002) such that the overall shift in autonomic balance (LF/HF) was not affected by m-CLD on day 1 (**Figure 3E**; effect of stress: *F_1,25_* = 4.77, *p* = 0.04) nor day 5 (**Figure 3I;** effect of stress: *F_1,23_* = 6.41, *p* = 0.02). Further examination of the autonomic response to WS exposure revealed that WS-evoked changes in blood pressure were primarily driven by changes in heart rate with little to no effect of m-CLD (see supplemental **Figure S10**).

### 3.6 Intra-LC m-CLD has no Effect on Sucrose Preference (Study C)

Repeated WS resulted in a modest reduction in sucrose preference among witnesses compared to controls (**Figure 3H**; effect of stress: *F_1,43_* = 8.05*; p* = 0.01; CON + vehicle, n=11; CON + m-CLD, n=11; WS + vehicle, n=14; WS + m-CLD, n=11). However, post hoc analyses revealed no significant differences, and intra-LC m-CLD therefore had no effect on this stress-induced reduction in sucrose preference (effect of treatment: *F*_1,43_ = 0.80, *p* = 0.38). Importantly, a pre-stress sucrose preference test performed prior to vehicle or m-CLD treatment confirmed that there were no pre-existing differences in sucrose preference between witnesses (n = 20) and controls (n = 18) prior to the start of stress manipulations (unpaired *t* test, *t*_36_ = 1.02, *p* = 0.31; data not shown).

### 3.7 Resting CORT Levels are Increased by a History of WS, but Unaffected by Intra-LC m-CLD (Study C)

While Study A demonstrated WS-induced increases in CORT in response to an acute stress exposure, Study C measured resting plasma CORT collected six days following the cessation of WS (**Figure 3K**), revealing that a history of WS (WS + vehicle, n = 6; WS + m-CLD, n = 5) elicited elevated resting levels of CORT as compared to controls (CON + vehicle, n = 5; CON + m-CLD, n = 4). (**Figure 3K**, effect of stress *F_1,16_* = 7.7 *p* = 0.01). However, intra-LC m-CLD did not impact this stress-induced increase in plasma CORT, (**Figure 3K**; effect of treatment: *F_1,16_* = 2.4, *p* = 0.14).

### 3.8 Intra-LC m-CLD Prevents Persistent WS-Induced Increases in IL-1β (Study C)

As measured at rest within the LC, persistent increases in IL-1β were identified within the LC in rats with a history of WS and this was blocked by prior treatment with m-CLD (**Figure 3L**, stress x drug interaction: *F_1,14_* = 11.47, *p* = 0.004; CON + vehicle (n = 5); CON + m-CLD (n = 5); WS + vehicle (n = 4); WS + m-CLD (n=4)). Notably, this accumulated neuroimmune response under resting conditions in rats with a history of WS was specific to IL-1β as there was no effect of WS nor m-CLD on any of the other 9 cytokines measured (**Table S1**).

### 3.9 Intra-CeA m-CLD had no effect on WS-evoked behavior (Study D)

Studies first confirmed that the 25μg dose of m-CLD injected into the CeA was equally as effective at ablating microglia as was identified in the LC (**Figure S8**). Unlike the robust behavioral effects observed following intra-LC m-CLD, microglial ablation in the CeA had no impact on WS-evoked behaviors. Burying was increased among WS rats compared to control rats on Day 1 as expected, and this was not affected by intra-CeA m-CLD (**Figure 4B**, effect of stress: *F_1,47_* = 53.9, *p* < 0.0001; effect of treatment: *F_1,47_* = 3.2, *p* = 0.082; CON + vehicle, n = 13; CON + m-CLD, n = 13; WS + vehicle, n = 12; WS + m-CLD, n = 13). Similar findings were observed on Day 5 (**Figure 4C**, effect of stress: *F_1,49_* = 76.3, *p* < 0.0001; effect of treatment: *F_1,49_* = 0.6, *p* = 0.5). Freezing behavior did not differ between WS rats treated with m-CLD and WS rats treated with vehicle on Day 1 (**Figure 4D**, WS + vehicle vs. WS + m-CLD unpaired *t* test, t_23_ = 1.41, *p* = 0.17) or Day 5 (**Figure 4E**, WS + vehicle vs. WS + m-CLD, unpaired *t* test, *t*_24_ = 0.31, *p* = 0.92). Rearing behavior was not affected by WS nor by intra-CeA m-CLD on day 1 (**Figure S11C**) or day 5 (**Figure S11D**).

### 3.10 Intra-CeA m-CLD Differentially Regulates Resting and WS-Evoked IL-1β (Study D)

In the present study, tissue collected from rats at rest five days following WS/control did not exhibit increased IL-1β in the CeA (**Figure 4F**, F_1,14_ = 3.5, p = 0.08). Previously, we reported that WS context exposure conducted 5 days after the final day of WS/CON produced an exaggerated IL-1β response in the CeA selectively among female rats with a history of WS exposure (Finnell et al., 2018). These data suggest that a subthreshold stress challenge (WS context) is necessary to reveal the neuroimmune sensitization that occurred in rats with a history of WS (unlike the LC). Therefore, a separate subset of rats was exposed to WS 60 minutes prior to tissue collection to determine whether intra-CeA m-CLD suppressed the WS-induced neuroinflammatory priming we previously observed (Finnell et al., 2018). Confirming our prior findings, rats with a history of WS exposure do indeed exhibit greater IL-1β in the CeA following WS context exposure, which is blocked by m-CLD (**Figure 4G**, stress history x drug interaction: F_1,22_ = 5.31, *p* = 0.031). Taken together, these data show that mCLD inhibited the WS-evoked neuroimmune response when administered into the CeA.

## 4. DISCUSSION

There is a clear clinical correlation between social stress and the development of anxiety and depression, with women at increased risk for these stress-related disorders (Kendler et al., 1999; Kessler, 2003; Sandanger et al., 2004). A strong body of literature supports a causal role for neuroimmune dysregulation in the pathophysiology of these conditions (Calcia et al., 2016; Finnell & Wood, 2018; Frank et al., 2016). However, preclinical studies examining region-specific effects of microglia in the pathophysiology of stress-induced psychosocial disorders, especially among females, are severely lacking. To our knowledge, this is the first study to establish a role for LC microglia in regulating the neuroimmune and hypervigilant behavioral responses to social stress in females. Specifically, these studies determined that microglial activation using the TLR4 agonist, LPS, within the LC exacerbates the hypervigilant-like burying behavior and peripheral inflammation in response to an acute social stressor. Further, reducing the number of microglia within the LC via discrete microinjection of m-CLD attenuates WS-evoked burying behavior, blocks WS-evoked vagal withdrawal, and prevents persistent WS-induced increases in inflammatory cytokines in the LC, with minimal impact when administered into the CeA.

Prior studies have demonstrated that minocycline effectively attenuates stress-induced behavioral deficits, highlighting a role for macrophages in stress susceptibility (Liu et al., 2018; Moller et al., 2016; Rooney et al., 2020; Soczynska et al., 2012). Further, CSF1R antagonists administered in the diet demonstrate a specific role for microglia (Lehmann et al., 2019), but these methods do not target inflammatory cells within specific brain regions. Recently, clodronate disodium salt has been used to induce transient microglial depletion without altering the expression of genes defining neurons or other immune cells within the brain (Schalbetter et al., 2022). Additionally, non-mannosylated formulations of liposomal CLD have been used in the brain to induce microglial depletion (Xie et al., 2017). However, contradictory evidence suggests low cell-type specificity of non-mannosylated liposomal CLD as another recent study reported that CLD caused neuronal damage when used in this non-specific manner (Han et al., 2019). Recent development of m-CLD, where CLD is encapsulated in mannosylated liposomes, allows CLD to be selectively taken up by microglia due to the microglial-specific expression of mannose receptors (Burudi & Regnier-Vigouroux, 2001). In the current study, we found no evidence of neuronal damage using this mannose-targeted liposomal formulation of CLD. Importantly, this study is the first to characterize the effects of m-CLD administered in discrete stress-sensitive brain regions, highlighting a novel method by which researchers can effectively target microglia in a site-specific and functional manner without inducing neuronal damage.

The LC is well-recognized to regulate hypervigilant burying behavior in the context of stress as chemical depletion of LC-NE tone inhibits CRF- and stress-evoked burying (Howard et al., 2008). Despite longstanding knowledge from electrophysiological studies that IL-1β activates LC neuronal firing through IL-1 receptor (Borsody & Weiss, 2002a, 2002b; Morris et al., 2020), the role of LC microglia in the context of social stress was unknown. Persistently increased LC activity is linked to anxiety (Morris et al., 2020), suggesting that stress-induced activation of LC microglia may promote anxiety disorders. Importantly, the current studies identified that intra-LC microglial knockdown prevents persistent WS-induced increases in IL-1β in the LC. While it has generally been assumed that LC stress responses were elicited via corticotropin-releasing factor (CRF) receptors on LC-NE neurons, these data provide the first evidence of a significant contribution of LC microglial cells towards LC dependent behaviors. Importantly, CRF receptors are also expressed on microglia, and future studies will be critical in determining if CRF-mediated microglial activation facilitates hypervigilant behavioral responses. The behavioral effects of microglial depletion were specific to the LC as intra-CeA m-CLD effectively reduced WS-evoked IL-1β with no effect on behavior. It is not surprising that intra-CeA m-CLD did not affect WS-evoked burying as the CeA does not directly regulate burying behavior. However, we also analyzed WS-evoked freezing, a behavioral response which is regulated by the CeA (Choi & Brown, 2003), and found that intra-CeA microglial depletion had no effect. However, we cannot rule out that there may be implications for this effect of m-CLD if other behavioral tests regulated by CeA were conducted. For example, reductions in WS-evoked neuroinflammation in the CeA may subsequently affect behavior during elevated plus maze or acoustic startle testing.

Notably, m-CLD does not permanently reduce microglia, and while it is unknown exactly how long the microglia are reduced by m-CLD in the present study, they have likely repopulated by the time the brains were collected as suggested by previous studies (Nelson & Lenz, 2017; Schalbetter et al., 2022; Torres et al., 2016; VanRyzin et al., 2016). In light of this consideration, it is striking that the functional effect of transient LC microglial knockdown is maintained, as evidenced by the blockade of accumulating IL-1β in the LC under resting conditions up to six days after the final stressor. It is also important to note that although the WS-evoked burying response is interpreted as a hypervigilant behavior, and LC microglia regulate this response during WS, we cannot conclude that this response is maladaptive in nature or perhaps signifies a coping response. Additional post-stressor behavioral assays such as acoustic startle or marble burying need to be conducted to better understand the implications of these findings in the greater context of the development of a persistent hypervigilant phenotype.

Beyond behavioral regulation, modulation of LC activity can regulate other aspects of the stress response (e.g., autonomic response and HPA axis activity) due to widespread LC-NE projections (Loughlin et al., 1986; Szabadi, 2013; Uematsu et al., 2015; Valentino & Van Bockstaele, 2008; Wood & Valentino, 2017). The autonomic response to stressors is a highly differentiated pattern of activity across a range of physiological measures, including but not limited to changes in HRV and blood pressure. Importantly, the LC-NE projections to the vagus facilitate vagal withdrawal (Samuels & Szabadi, 2008; Wang et al., 2014; Wood & Valentino, 2017). Here, we found that CLD had no overall effect on WS-evoked autonomic balance as measured by LF/HF ratio of HRV. However, a more detailed inspection of vagal tone, indicated by the HF component of HRV, revealed that repeated WS resulted in vagal withdrawal, and LC microglial ablation blocked this response. Therefore, a compensatory sympathetic response, independent of LC microglia, appears to co-regulate autonomic function during repeated WS exposure. The compensatory increase in sympathetic drive also likely explains the increased MAP observed during WS as we demonstrated that this effect was primarily driven by increased heart rate, another indication of sympathetic dominance. Nonetheless, these are the first data to suggest that LC microglia facilitate a reduction in vagal tone in response to repeated social stress exposure. It is also noteworthy that m-CLD-treated controls do not exhibit as clear of a habituation, in terms of the hemodynamic response, to the handling as vehicle-treated controls, suggesting that microglia within the LC may play distinct regulatory roles in the context of WS compared to control, and future studies should aim to better understand this dichotomous role for LC microglia.

Based on our current findings, m-CLD blunted the dynamic response in control rats but had little effect on the hemodynamic responses induced by WS, further supporting the idea that increased sympathetic drive independent of LC microglia counteracts the effects of m-CLD on autonomic function. Interestingly, this compensatory increase in sympathetic drive that occurred in m-CLD-WS rats was specific to the cardiovascular arm of the autonomic response, as WS-evoked hypervigilant behavior was persistently suppressed by m-CLD. Future studies are needed to explicate the effects of LC microglia on the dynamic cardiovascular responses in both the control and WS animals. Moreover, emerging evidence suggests that microglia-neuron interactions vary depending on the neuronal population (Bolton et al., 2022; Favuzzi et al., 2021; Rosin et al., 2021). Thus, additional studies are needed to characterize LC microglia-neuron interactions and their implications for differential regulation of the stress response.

Here, we also show that plasma CORT is increased immediately following WS exposure, and that intra-LC LPS alone produces a comparable CORT response in non-stressed rats. This is not surprising given the stimulatory role that the LC has on the HPA axis. Further, these studies found that rats with a history of repeated WS exposure exhibit persistently elevated resting CORT levels independent of prior treatment with m-CLD. Thus, the current findings indicate that intra-LC microglia may facilitate acute HPA activation, but do not impact the persistent increases that are consistent with repeated stress exposure and many psychiatric disorders. It is noteworthy that the resting CORT levels of the control rats are slightly higher than typical basal physiological conditions (Windle et al., 1998) which is likely due to repeated handling exposure and the isolation stress of being singly housed. Additionally, sucrose preference was significantly reduced among females repeatedly exposed to WS. However, this response was moderate and not affected by LC microglial manipulation. The LC is not traditionally considered to regulate anhedonia, so these findings are in line with the selectivity of ablating LC microglia rather than whole brain.

These studies have provided a significant advance in our understanding of how microglia may function to regulate behavior in the LC. However, it is also important to acknowledge the limitations of these studies. First, LPS is a TLR4 agonist, but is not selective for microglia, and therefore other cells, including neurons, astrocytes, and endothelial cells may contribute to the intra-LC LPS-induced burying responses observed. Further, upon assessing peripheral IL-1b, we found that intra-LC LPS does cause an increase, but that WS + LPS causes an even greater increase, which parallels the behavioral findings presented in this study. Importantly this peripheral immune response itself did not affect anxiety-like behavior as controls treated with LPS did not bury more than vehicle controls (Figure 1B). Additionally, because LC neuronal activity was not measured in these studies, we cannot make direct conclusions about the impact of microglial expression on LC neuronal activity and highlights an important future direction. Finally, because our analysis was focused on the potential compensatory effects that were possible in the context of WS that may counteract the effect of microglial depletion, we do not have a measure of HMGB1 or CX3CL1 among control-handled rats treated with m-CLD or vehicle, so we cannot draw conclusions about the effect of m-CLD on these endpoints in the absence of WS exposure.

## 5. CONCLUSIONS

Overall, these studies examined the role of LC microglia in response to acute and repeated WS exposure in females in two ways: first, by activating microglia with a low dose of LPS and second, by reducing the number of microglia in the LC with m-CLD. Together, these studies identify a clear role for LC microglia in regulating anxiety-like hypervigilant behavior induced by WS. In characterizing the effects of intra-LC m-CLD treatment before, during, and after WS exposure, these studies establish an effective way to selectively target microglia in a region-specific manner. The current results these data provide the first evidence of a significant contribution of LC microglial cells towards regulating behavioral and physiological responses to psychosocial stress. In future studies, it should be determined if suppression of hypervigilant responses during social stress by microglial knockdown, corresponds to changes in long-term susceptibility to hypervigilance. However, the robust and lasting suppression of IL-1β suggests that modulation of microglia may have lasting protective effects in the context of social stress.

## 6. ACKNOWLEDGEMENTS

Hippocampal tissue used in supplementary studies was generously donated by our collaborator, Dr. Lawrence Reagan (University of South Carolina School of Medicine). This work was supported by the National Institutes of Health (R01 MH113892) and Veterans Health Administration (I21 BX002664).

## DISCLOSURES

The authors have no conflicts of interest to disclose.

## Notes

### Competing Interest Statement

The authors have declared no competing interest.

