## Supplemental Info for "Site-Specific Knockdown of Microglia in the Locus Coeruleus Regulates Hypervigilant Responses to Social Stress in Female Rats"

### Supplemental Methods

#### Characterization of m-CLD

*m-CLD Infusion, Dosage, and Drug Spread*

Previous studies aided in the dosage determinations tested in the present study. To our knowledge, only a few studies have been published that administered liposome-encapsulated CLD in the brains of rats (Drabek et al., 2012; Faustino et al., 2011; Han et al., 2019; Xie et al., 2017), however it should be noted that these were not mannoslyated formulations as was used here, and for several of the papers, the concentration of CLD was not provided, but rather only the infusion volume. The infusion volume administered intracranially ranged from 3µL into the right cortex of postnatal day 5 rats (Faustino et al., 2011) to 5µL into the hippocampus (Drabek et al., 2012) or 10µL into the substantia nigra (Xie et al., 2017) of adult rats. Importantly, Drabek et al., measured intracranial pressure during the 5µL administered over 10 minutes and showed no changes in intracranial pressure (Drabek et al., 2012). These studies also identified that liposome-encapsulated CLD was capable of reducing microglia by approximately 70% in the 24 hours following infusion (Faustino et al., 2011; Xie et al., 2017) while others confirmed continued reduction of 40% through day 7 (Drabek et al., 2012). It is important to note that microglial depletion is temporary and has been shown to fully repopulate by 14 (Bruttger et al., 2015) to 20 days (Torres et al., 2016) after ablation. Therefore, it is expected that microglia are beginning to repopulate throughout the duration of the current study design. The concentration of the m-CLD formulation used in the present study was 5 µg/µL. Rats were maintained under anesthesia during the infusion (1-2% isoflurane). Infusions (m-CLD and empty mannoslyated liposomes (vehicle) were administered via the indwelling cannulas aimed at the LC or CeA and 5uL was administered over a 15-minute period. Infusions were conducted 3 days prior to the first day of WS or control.

To confirm that the slow infusion of 5µL of m-CLD over 15 minutes did not spread beyond the LC region, we infused a fluorescent mannoslyated liposome (Encapsula Nanosciences, CLD-8929) directly into the LC (**Figure S4A**). The size of the LC was determined using Dopamine-β hydroxylase (DβH) immunofluorescence (anti-mouse DβH, 1:3000, Millipore MAB308) and anti-mouse Alexa Fluor 546 (1:1000, Thermofisher A11030, **Figure S4B**).

*Brain Tissue Perfusion*

Rats were transcardially perfused with 4% paraformaldehyde, dissected and sectioned into an anterior and posterior section, and then immersed in 20% sucrose with 1% sodium azide until saturated. Upon saturation, brains were snap-frozen in ice-cold 2-methylbutane and stored at -80°C until further processing.

*Immunohistochemistry for Fluoro-Jade B and Iba-1*

Slides were washed in nanopure water (Millipore) and incubated in 0.001% Fluoro-Jade B in acetic acid and nanopure water (Millipore) at room temperature for 30 minutes followed by a final rinse in nanopore water. Importantly, Fluoro-jadeB has also been identified as a marker for activated microglia (Damjanac et al., 2007), and therefore slices were double labeled for Iba-1 (1:1000, overnight at 4°C, Wako) followed by donkey anti-rabbit fluorescent secondary (1:200, Fisher Dylight 594). Slides were cover-slipped using Fluoromount-G (Southern Biotech, Birmingham, AL) prior to assessment.

Neuronal damage was quantified as the total number of Fluoro-Jade B positive cells (i.e., all damaged neurons and activated microglia, shown in green below, **Figure S3A**) minus the number co-labeled Fluoro-Jade B and Iba-1 (only activated microglia, red not shown). Hippocampal tissue from an adrenalectomized rat was generously donated by Dr. Lawrence Reagan (University of South Carolina School of Medicine) as a positive control due to the significant neuronal death that occurs in the hippocampus following adrenalectomy (**Figure S3B**).

*Immunohistochemistry for Iba-1 in the CeA:*

CeA microglial knockdown validation: To verify microglial knockdown in Study D, Iba-1, was quantified in brains treated with 25μg m-CLD or vehicle (empty liposomes). Free floating sections were washed in 0.1M PBS followed by 0.75% hydrogen peroxide in 0.1M PBS to block endogenous peroxidases. Tissue was then washed in 0.1M PBS before blocking in 3% normal goat serum (NGS) in 0.1M PBS + Triton X-100 for 60 minutes. Slices were incubated overnight at 4°C in the primary antibody rabbit anti-rat Iba-1 (1:1000, Wako Pure Chemical Industries, Richmond, VA) diluted in 0.1M PBS+1% NGS. Tissue was washed in 0.1M PBS and incubated in the secondary antibody, biotinylated goat anti-rabbit IgG (Vector Laboratories, Burlingam, CA), at room temperature for 90 minutes. Following three 5-minute washes in 0.1M PBS, tissue was incubated in an avidin-biotin complex (Vector Laboratories) for 30 minutes at room temperature before a final wash in 0.1M PBS. Tissue was then processed with metal-enhanced diaminobenzidine tetrahydrochloride (DAB) (Sigma-Aldrich) for the same incubation time, washed, mounted onto slides and cover slipped. Tissue was imaged and the number of Iba-1 positive cells in the CeA was quantified using a region of interest that was kept constant for all slices. See **Figure S4**.

*Cardiovascular Measures: Cardiac Output, Total Peripheral Resistance and Stroke Volume*

In addition to changes in brain signaling, immune function, and behavior, social stress also results in shifts in autonomic function, often marked by changes in cardiovascular function. These differentiated patterns represent an integrated response to environmental challenges that vary as a function of type of challenge, characteristics of the organisms responding, the opportunity or lack thereof of a coping response, and may vary over time. Winters et al., succinctly described two common responses patterns, the defense reaction and the vigilance reaction (Winters et al., 2000). The autonomic cardiovascular response pattern associated with the defense reaction is characterized by elevations in blood pressure that are due to elevated cardiac output (CO) whereas the vigilance reaction is characterized by elevations in blood pressure due to elevated total peripheral resistance (TPR). In addition, the defense reaction is more likely to occur when an organism has an opportunity to emit a coping response such as fight or flight (active coping) whereas the vigilance reaction is more likely to occur when such an opportunity to respond is less likely (passive coping). Importantly, when exposed to a stressor the elevation in blood pressure may be initially due to elevations in CO but over time (both within session and between sessions) may transition to be supported by elevations in TPR (Ring et al 2002). This may represent an appraisal by the organism that the likelihood of an effective active coping response has decreased and thus a passive coping response is more appropriate. To further investigate the vigilance response in stress-exposed female rats we used telemetry data collected in Study C to measure mean arterial blood pressure (MAP). Using these measurements, we calculated the underlying hemodynamics of cardiac output (CO) and total peripheral resistance (TPR; MAP= CO x TPR) (See Figure S9) based on validated equations (Hill et al., 2012, 2013; Sun et al., 2005). Since CO is a function of stroke volume (SV) and heart rate (HR) (CO=SV x HR), we further probed the CO response by examining SV and HR.

### Supplemental Figures

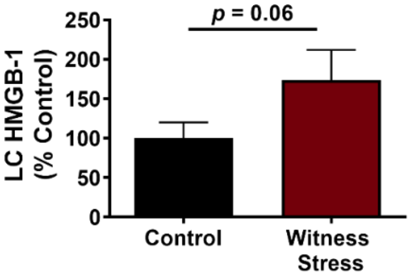


**Figure S1 – Repeated witness stress (WS) produces accumulation of HMGB-1 in the LC.** Six days after the 5th (final) day of WS or control handling exposure, HMGB-1 is increased in the LC of witnesses (n=6; 173.9 ± 38.0) compared to controls (n=5; 100.0 ± 20.1). Data presented as % control (unpaired *t* test $t_{7.5} = 1.7$, $p = 0.06$).

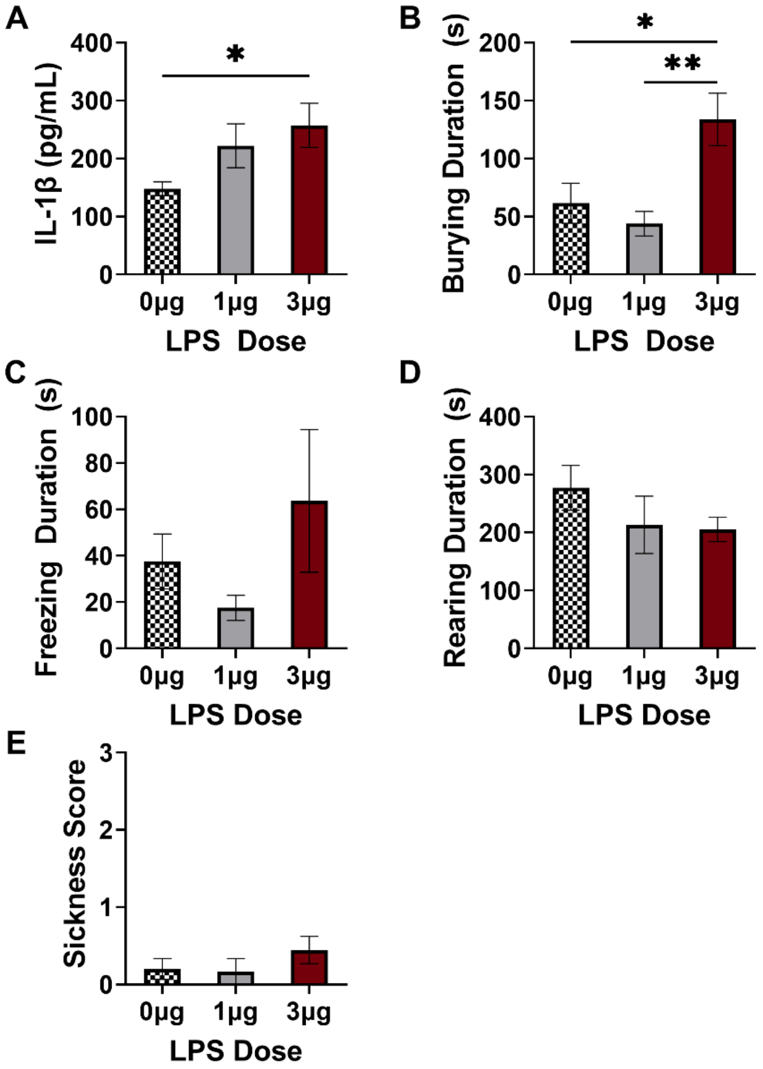


**Figure S2**. **Intra-LC lipopolysaccharide (LPS) dose response.** Plasma levels of IL-1β increases in a dose-dependent manner with the 3μg dose of LPS inducing the greatest plasma IL-1β response ($F_{2,12} = 0.6923$, $p = 0.044$; *$p < 0.05$ 0μg vs. 3μg). The 3μg dose of intra-LC LPS also induced significantly greater burying during WS compared to vehicle (*$p < 0.05$) and 1μg (**$p < 0.01$). Freezing ($F_{2,18} = 1.548$, $p = 0.240$) and rearing ($F_{2,20} = 1.298$, $p = 0.295$) behavior during WS were not affected by pretreatment with intra-LC LPS. Importantly, the 3μg dose did not induce sickness behavior (**E**) compared to vehicle ($p = 0.497$) or 1μg LPS ($p = 0.497$).

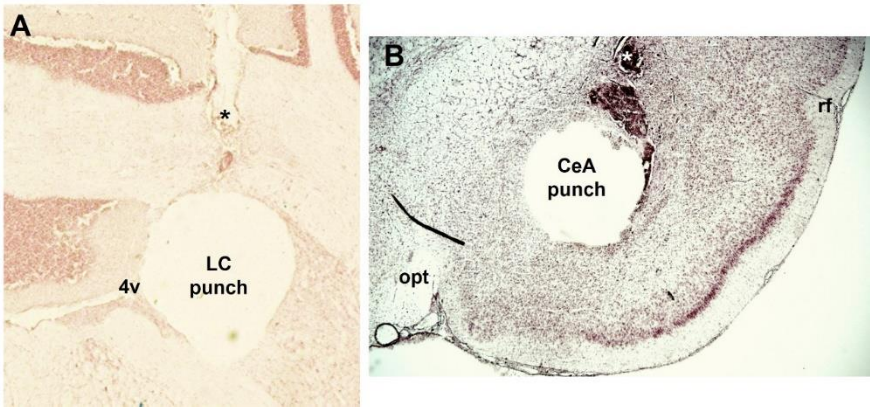


**Figure S3. Histological verification of cannula and punch placement.** Tissue slices (30 μM) were collected before and after tissue collection, stained with neutral red and assessed using a microscope. Representative images of the locus coeruleus (LC; **A**) and central nucleus of the amygdala (CeA; **B**) tissue punch and cannula placements are shown above. ***** denotes cannula track, rf = rhinal fissure, opt = optic tract, 4v = 4$^{th}$ ventricle.

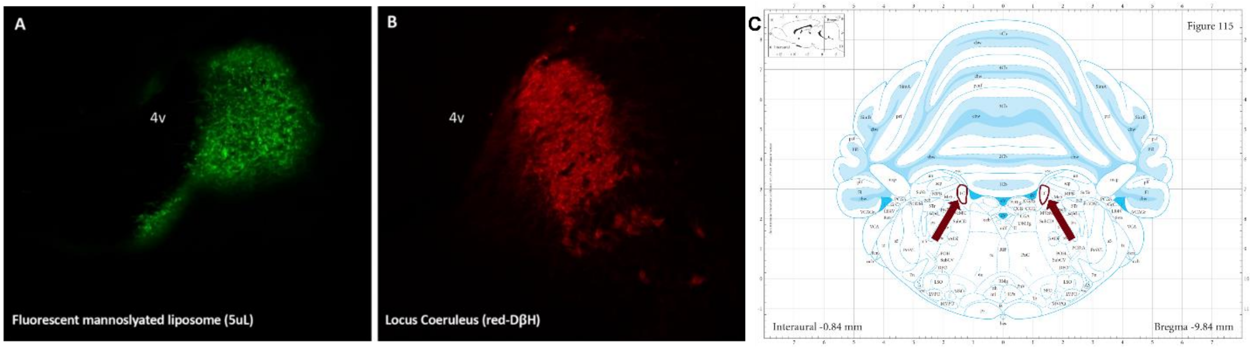


**Figure S4**. **Intra-LC fluorescent mannosylated liposomes confirm spread of drug does not expand beyond the LC.** A representative image of the infusion spread (**A**) and LC DβH (**B**) are shown above. An average of three separate rat's LC indicates that the average area of the core of LC is average ± SEM: 156,096±14,161 pixels$^2$ (Image J, NIH). The infusion spread was determined using injections of 5μL of fluorescent liposomes into the LC region in two separate rats and found to be contained within an average of 151,303 ± 7,513 pixels$^2$ (Image J, NIH). Importantly, because the LC is positioned directly next to the 4$^{th}$ ventricle (**C**), the rostral caudal extent of brains from the fluorescent liposome injected rats were evaluated for evidence of fluorescent dye by collecting every 4$^{th}$ slice throughout the brain. Fluorescent liposomes were not identified in any brain region outside the LC.

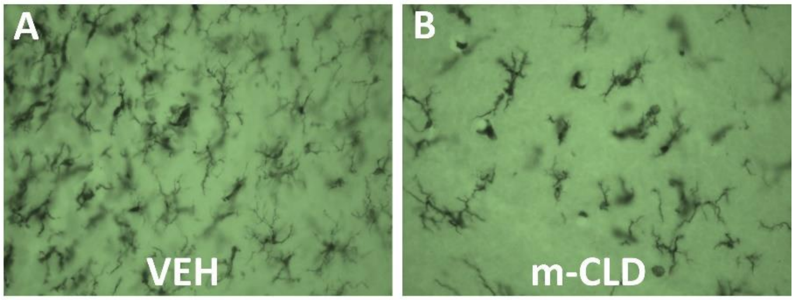


**Figure S5. m-CLD effectively reduces microglial expression within the locus coeruleus (LC).** Three days following intra-LC empty liposomal vehicle (**A**; 0µg) or m-CLD (**B**; 25µg), immunohistochemical detection of Iba-1 confirmed microglial expression was reduced by approximately 50% in agreement with the findings presented in the main text (**Figure 2B**).

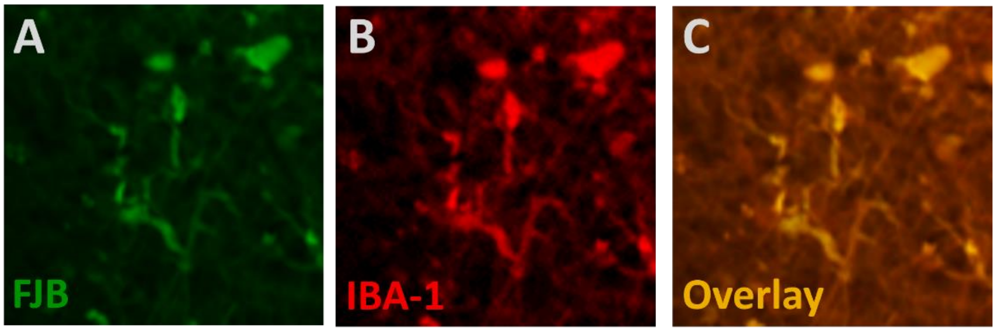


**Figure S6 – m-CLD does not induce neuronal damage.** As an extension of figure 2C, histological staining using FJB has been shown to stain both degenerating neurons and activated microglia as discussed in the main text. Therefore, slices were stained for both Fluorojade-B (FJB) and IBA-1 and only FJB positive/IBA-1 negative cells were counted as dying neurons. A representative cluster of FJB positive **(A; green)** and IBA-1 positive **(B; red)** cells are depicted, and an overlay of the two images is shown in panel **C**.

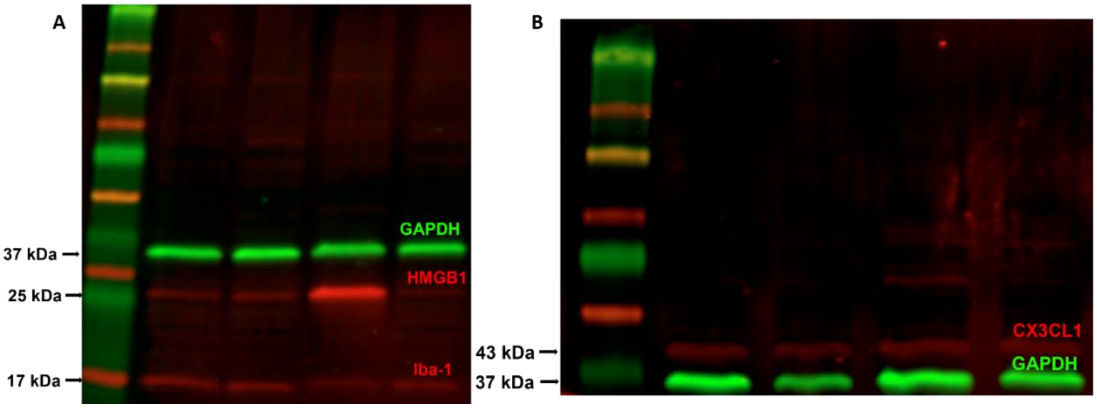


**Figure S7 – Representative Western Blot Images.** A representative image for a western blot depicting HMGB-1 at 25 kDa, Iba-1 at 17 kDa, and our housekeeping protein, GAPDH at 37 kDa is depicted on the left (**A**). On the right (**B**), another blot shows detection of CX3CL1 in red at 43 kDa with our housekeeping protein, GAPDH, in green at 37 kDa.

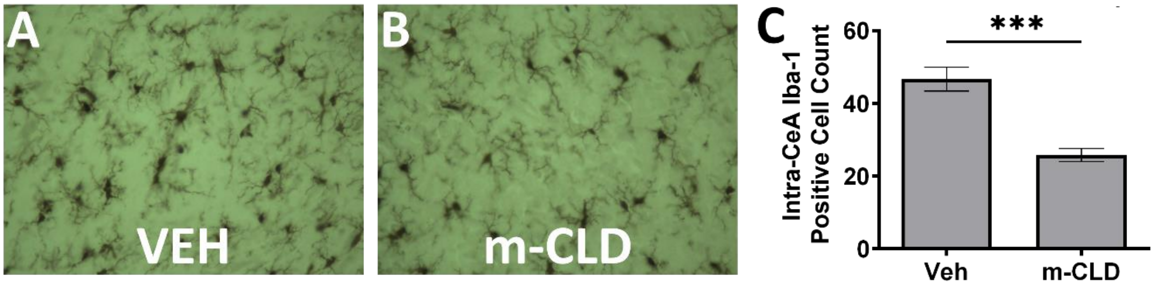


**Figure S8 – m-CLD effectively reduces microglial expression within the CeA.** Three days following microinjection of intra-CeA empty liposomal vehicle (**A**; 0µg) or m-CLD (**B**; 25µg), immunohistochemical detection of Iba-1 confirmed microglial expression was reduced by approximately 50% (**C**; Paired *t* test, $t_7 = 6.793$, $p = 0.0003$; data shown as mean ± SEM).

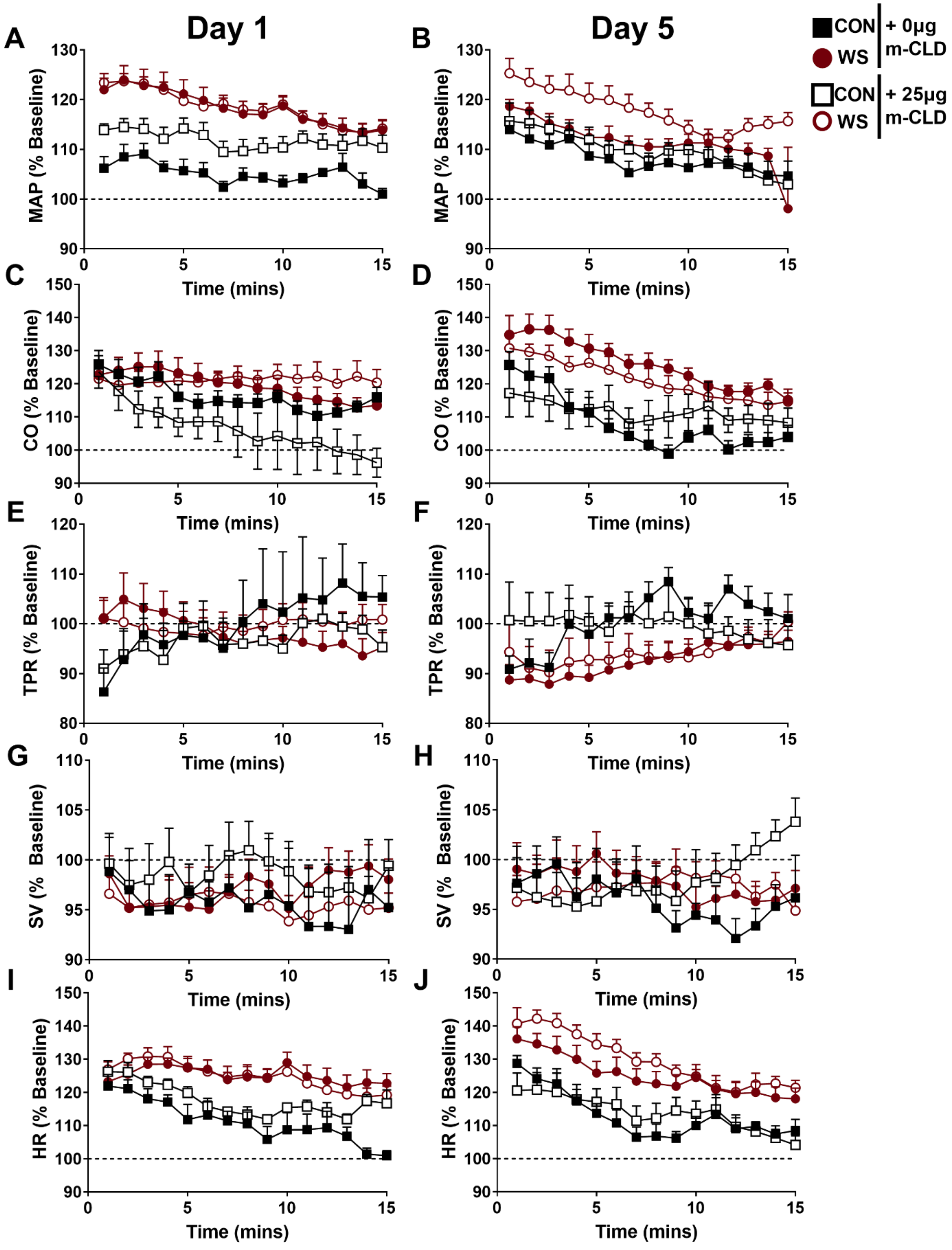


**Figure S9. Hemodynamic responses during WS/CON exposure on Day 1 and Day 5.** MAP was elevated to a greater extent among WS rats compared to controls on both

Day 1 (**A**; effect of stress: $F_{1,24} = 23.81$, $p < 0.0001$) and Day 5 (**B**; effect of stress: $F_{1,23} = 8.51$, $p = 0.01$). Despite no effect of m-CLD on Day 1 (effect of drug: $F_{1,24} = 2.73$, $p = 0.11$), on Day 5, a noteworthy trend was observed in which m-CLD appears to enhance MAP to a greater extent in response to WS (**B**; effect of drug: $F_{1,23} = 4.10$, $p = 0.05$). WS-induced elevations in MAP were accompanied by elevated CO on Day 1 (**C**; effect of stress: $F_{1,24} = 7.48$, $p = 0.02$) and Day 5 (**D**; effect of stress: $F_{1,23} = 13.86$, $p = 0.001$). TPR was not impacted by WS on Day 1 (**E**; effect of stress: $F_{1,24} = 0.05$, $p = 0.83$), but following repeated WS, on Day 5, TPR was reduced among WS rats (**F**; effect of stress: $F_{1,23} = 4.49$, $p = 0.04$) compared to controls, which may indicate that with repeated WS exposure, female rats shift from a vigilant hemodynamic response to a more defensive hemodynamic response. However, there was no effect of m-CLD on TPR on Day 1 (**E**; effect of drug: $F_{1,24} = 0.08$, $p = 0.78$) nor following repeated WS exposure (**F**; effect of drug: $F_{1,23} = 0.01$, $p = 0.91$). SV was changed little during WS exposure regardless of m-CLD treatment on Day 1 (**G**; effect of stress: $F_{1,24} = 0.11$, $p = 0.75$) as well as Day 5 (**H**; effect of stress: $F_{1,24} = 0.08$, $p = 0.78$). Thus, not surprisingly, the WS-evoked increases in CO is primarily driven by elevated HR among WS rats on both Day 1 (**I**; effect of stress: $F_{1,24} = 19.85$, $p = 0.0002$) and Day 5 (**J**; effect of stress: $F_{1,23} = 18.50$, $p = 0.0003$).

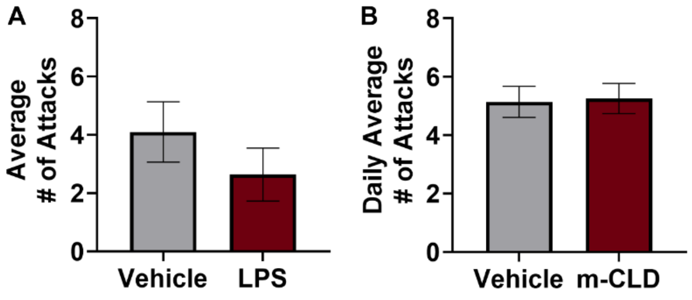


**Figure S10. Average number of attacks observed during WS.** The average number of attacks observed by witnesses in study A did not differ by drug treatment (**A**; mean ± SEM: vehicle WS, 4.1 ± 1.04; LPS WS, 2.6 ± 0.91; unpaired *t*-test $t_{19} = 1.07$, $p = 0.30$). Similarly, the average number of attacks observed across all 5 days of WS in Study C did not differ by drug treatment (**B**; mean ± SEM: WS + VEH, 5.15 ± 0.0.53; WS + m-CLD, 5.26 ± 0.52; unpaired *t* test: $t_8 = 0.15$, $p = 0.88$).

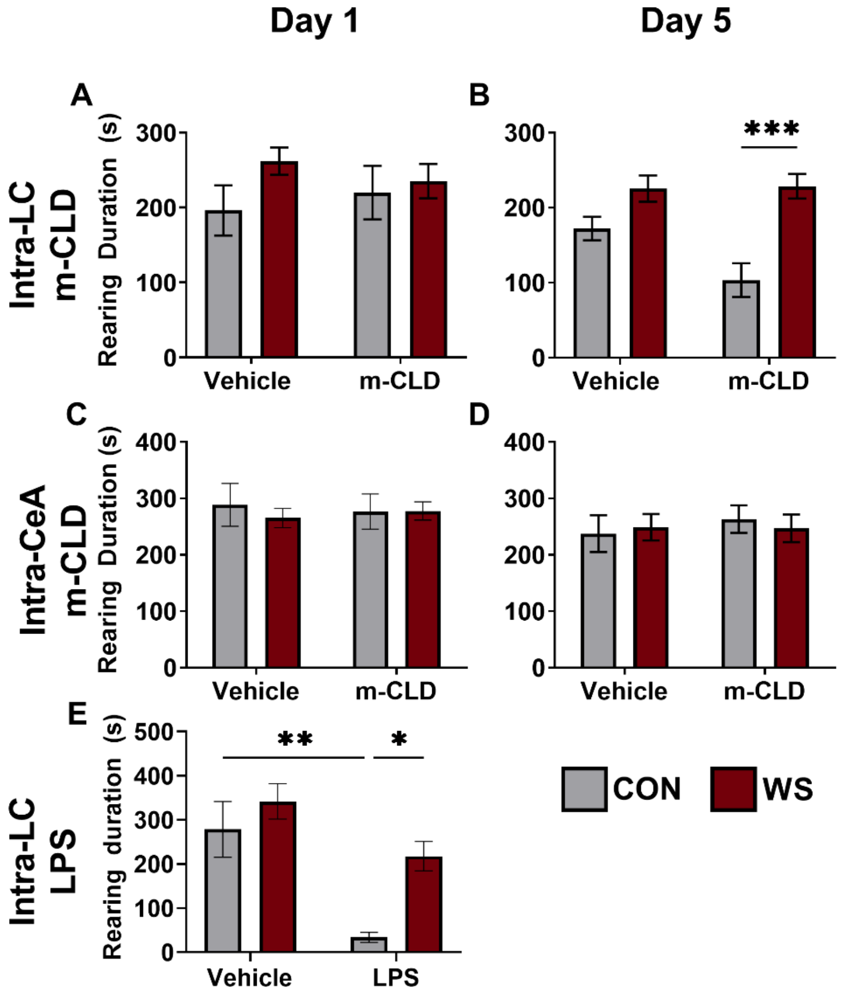


**Figure S11. Rearing behavior during WS and CON exposure.** In Study C (**A, B**), there was no effect of WS nor intra-LC m-CLD treatment during day one (**A**; effect of stress: $F_{1,46}$ = 2.05, $p$ = 0.16; effect of drug: $F_{1,46}$ = 0.002, $p$ =0.10; stress x drug interaction: $F_{1,46}$ = 0.79, $p$ = 0.38). However, during day 5, within m-CLD-treated rats, rearing was significantly reduced among controls compared to witnesses (**B**; CON + m-CLD vs. WS + m-CLD, ***$p$ = 0.0008). There was no effect of WS nor m-CLD on rearing behavior in Study D on day 1 (**C**) or day 5 (**D**), suggesting that intra-CeA m-CLD treatment has no effect on rearing behavior during witness stress or control. Interestingly, similar to intra-LC m-CLD on day 5, intra-LC LPS (Study A) also results in a reduction in rearing behavior among controls rats (CON + LPS vs WS + LPS *$p$ = 0.03; CON + VEH vs CON + LPS, **$p$ = 0.004).

### Supplemental Tables

**Table S1. Cytokine expression in LC tissue collected at rest 6 days following the final witness stress/control exposure.** Data are expressed as percent of vehicle-treated control (mean ± SEM) for each analyte. Cytokine expression for vehicle-treated controls were as follows: IL-1β 19.7±4.8 pg/mL, IL-6 480.8±51.8 pg/mL, IL-10 27.3±0.9 pg/mL, IL-13 48.4±4.5, IFN-γ 38.7±10.6, MCP-1 17.7 ± 4.1 pg/mL IL-2, IL-4, GM-CSF and TNF-∝ were not detectable in these samples. *p<0.05 vs. control + veh. #p<0.05 vs witness + veh.

| Bioplex Analyte | Vehicle Treatment | | Clodronate Treatment | | Statistical Analyses | | |
|---|---|---|---|---|---|---|---|
| | Control | Witness | Control | Witness | Stress | Treatment | Stress x Treatment Interaction |
| IL-1β | 100.0±24.2 | 275.3±54.5* | 155.5±29.9 | 91.1±32.4# | $F_{(1,14)}$=2.46, p=0.139 | $F_{(1,14)}$=3.32, p=0.09 | $F_{(1,14)}$=11.50, p=0.004 |
| IL-6 | 100.0±4.8 | 86.5±9.9 | 96.0±2.6 | 100.0±4.0 | $F_{(1,15)}$=0.59, p=0.454 | $F_{(1,15)}$=0.58, p=0.458 | $F_{(1,15)}$=1.94, p=0.184 |
| IL-10 | 100.0±1.4 | 92.9±6.7 | 103.3±4.2 | 102.5±4.3 | $F_{(1,15)}$=0.727, p=0.407 | $F_{(1,15)}$=1.93, p=0.186 | $F_{(1,15)}$=0.464, p=0.506 |
| IL-13 | 100.0±4.2 | 89.8±6.0 | 98.8±2.3 | 100.7±3.9 | $F_{(1,15)}$=0.90, p=0.358 | $F_{(1,15)}$=1.21, p=0.289 | $F_{(1,15)}$=1.91, p=0.188 |
| IFN-γ | 100.0±12.3 | 83.2±16.5 | 93.7±4.0 | 108.2±8.7 | $F_{(1,15)}$=0.01, p=0.923 | $F_{(1,15)}$=0.641, p=0.436 | $F_{(1,15)}$=1.78, p=0.202 |
| MCP-1 | 100.0±10.2 | 123.3±30.6 | 99.3±15.2 | 115.7±26.2 | $F_{(1,15)}$=0.824, p=0.378 | $F_{(1,15)}$=0.037, p=0.851 | $F_{(1,15)}$=0.025, p=0.877 |
| IL-2 | not detected | | not detected | | not detected | | |
| IL-4 | not detected | | not detected | | not detected | | |
| GM-CSF | not detected | | not detected | | not detected | | |
| TNF-α | not detected | | not detected | | not detected | | |

**Table S2. Cytokine expression in CeA tissue collected at rest 6 days following the final witness stress/control exposure.** Data are expressed as percent of vehicle-treated control (mean ± SEM) for each analyte. Cytokine expression for vehicle-treated controls were as follows: IL-1β 70.3±33.3 pg/mL, IL-6 213.2±35.4 pg/mL, IL-10 28.3±2.8 pg/mL, IL-13 52.3±5.2, IFN-γ 49.3±6.9 pg/mL, MCP-1 30.3±10.0 pg/mL

| Bioplex Analyte | Vehicle Treatment | | Clodronate Treatment | | Statistical Analyses | | |
|---|---|---|---|---|---|---|---|
| | Control | Witness | Control | Witness | Stress | Treatment | Stress x Treatment Interaction |
| IL-1β | 100±21.2 | 99.7±15.5 | 130.8±24.8 | 64.0±7.2 | $F_{(1,14)}$=3.46, p=0.084 | $F_{(1,14)}$=0.019, p=0.893 | $F_{(1,14)}$=3.40, p=0.087 |
| IL-6 | 100.0±3.1 | 93.8±5.7 | 82.9±14.8 | 90.4±4.2 | $F_{(1,14)}$=0.006, p=0.938 | $F_{(1,14)}$=1.84, p=0.197 | $F_{(1,14)}$=0.819, p=0.381 |
| IL-10 | 100.0±4.5 | 97.4±4.2 | 94.4±4.0 | 90.3±6.7 | $F_{(1,14)}$=0.406, p=0.534 | $F_{(1,14)}$=1.46, p=0.248 | $F_{(1,14)}$=0.019, p=0.893 |
| IL-13 | 100.0±4.4 | 95.8±6.4 | 95.6±5.8 | 89.0±4.0 | $F_{(1,13)}$=1.10, p=0.313 | $F_{(1,13)}$=1.18, p=0.297 | $F_{(1,13)}$=0.058, p=0.814 |
| IFN-γ | 100.0±6.2 | 83.9±9.4 | 75.0±19.7 | 80.4±8.1 | $F_{(1,14)}$=0.228, p=0.641 | $F_{(1,14)}$=1.63, p=0.223 | $F_{(1,14)}$=0.915, p=0.355 |
| MCP-1 | 100.0±14.8 | 242.1±93.0 | 277.4±72.6 | 164.3±60.9 | $F_{(1,14)}$=0.054, p=0.82 | $F_{(1,14)}$=0.629, p=0.441 | $F_{(1,14)}$=4.13, p=0.062 |
| IL-2 | not detected | | not detected | | not detected | | |
| IL-4 | not detected | | not detected | | not detected | | |
| GM-CSF | not detected | | not detected | | not detected | | |
| TNF-α | not detected | | not detected | | not detected | | |

**Table S3. Cytokine expression in CeA tissue collected following a witness-stress challenge 6 days after the final witness stress/control exposure.** Data are expressed as percent of vehicle-treated control (mean ± SEM) for each analyte. Cytokine expression for vehicle-treated controls were as follows: IL-1β 8.1±1.7 pg/mL, IL-2 410.1±18.9 pg/mL, IL-4 8.7±0.3 pg/mL, IL-6 124.3±4.3 pg/mL, IL-10 31.4±1.5 pg/mL, IL-13 104.9±3.1 pg/mL, GM-CSF 2.9±0.3 pg/mL, IFN-γ 306.7±12.9 pg/mL, TNF-α 7.6±0.7 pg/mL, MCP-1 9.7±0.6 pg/mL **p<0.01 vs control + veh. $^{\$}$p<0.05 vs stress + veh. $^{\#}$p<0.05 vs control + veh

| Bioplex Analyte | Vehicle Treatment | | Clodronate Treatment | | Statistical Analyses | | |
|---|---|---|---|---|---|---|---|
| | Control | Witness | Control | Witness | Stress | Treatment | Stress x Treatment Interaction |
| IL-1β | 100±20.8 | 244.1±33.9** | 174.6±49.3 | 135.0±20.0$^{\$}$ | $F_{(1,28)}$=3.26, p=0.082 | $F_{(1,28)}$=0.357, p=0.555 | $F_{(1,28)}$=10.1, p=0.004 |
| IL-2 | 100±3.4 | 101.7±2.7 | 95.0±4.9 | 90.8±1.8$^{\$}$ | $F_{(1,27)}$=0.181, p=0.674 | $F_{(1,27)}$=6.96, p=0.014 | $F_{(1,27)}$=0.954, p=0.338 |
| IL-4 | 100.0±3.4 | 99.3±3.5 | 79.4±7.4$^{\#\$}$ | 80.5±3.6**$^{\$}$ | $F_{(1,29)}$=0.002, p=0.962 | $F_{(1,29)}$=20.0, p<0.0001 | $F_{(1,29)}$=0.039, p=0.844 |
| IL-6 | 100.0±3.4 | 91.4±5.6 | 73.1±5.9** | 74.1±3.0**$^{\$}$ | $F_{(1,29)}$=0.765, p=0.389 | $F_{(1,29)}$=25.8, p<0.0001 | $F_{(1,29)}$=1.21, p=0.280 |
| IL-10 | 100.0±4.8 | 93.5±5.8 | 79.6±5.5$^{\#}$ | 79.5±2.5** | $F_{(1,29)}$=0.518, p=0.477 | $F_{(1,29)}$=14.3, p=0.0007 | $F_{(1,29)}$=0.495, p=0.488 |
| IL-13 | 100.0±3.0 | 94.4±5.4 | 77.5±4.6** | 78.0±2.7**$^{\$}$ | $F_{(1,29)}$=0.436, p=0.514 | $F_{(1,29)}$=25.1, p<0.0001 | $F_{(1,29)}$=0.608, p=0.442 |
| GM-CSF | 100.0±10.6 | 204.5±68.2 | 71.2±7.6 | 71.8±6.2 | $F_{(1,17)}$=1.56, p=0.229 | $F_{(1,17)}$=3.68, p=0.072 | $F_{(1,17)}$=1.53, p=0.234 |
| IFN-γ | 100.0±4.2 | 94.0±4.6 | 78.1±4.7$^{\#}$ | 76.1±3.1**$^{\$}$ | $F_{(1,29)}$=0.88, p=0.356 | $F_{(1,29)}$=22.1, p<0.0001 | $F_{(1,29)}$=0.226, p=0.638 |
| TNF-α | 100.0±9.0 | 110.3±20.1 | 76.6±11.5 | 62.4±7.7$^{\$}$ | $F_{(1,25)}$=0.02, p=0.889 | $F_{(1,25)}$=6.89 p=0.015 | $F_{(1,25)}$=0.814, p=0.376 |
| MCP-1 | 100.0±4.0 | 190.4±44.2 | 377.8±186.3 | 252.8±46.8 | $F_{(1,27)}$=0.057, p=0.813 | $F_{(1,27)}$=5.49, p=0.027 | $F_{(1,27)}$=2.20, p=0.149 |

**Table S4. Summary of Statistical Statements for Behavior, CORT, and HRV Measures in Study C.** Statistical analyses for effects of intra-LC m-CLD are summarized in the table below. All data below were analyzed via 2-way ANOVA. Significant main effects are indicated in **bold** while significant post hoc analyses are denoted by the following: $^{*}$*p* < 0.05 vs drug-matched control; $^{+}$*p* < 0.05 vs stress condition-matched control

| Outcome of Interest | Vehicle Treatment | | Clodronate Treatment | | Statistical Analyses | | |
|---|---|---|---|---|---|---|---|
| | Control | Witness | Control | Witness | Stress | Treatment | Stress x Treatment Interaction |
| Day 1 Burying | 9.2 ± 2.4 | 79.4 ± 8.3$^{*+}$ | 10.5 ± 3.3 | 27.0 ± 7.1$^{+}$ | **$F_{(1,39)}$ = 55.6, p < 0.0001** | **$F_{(1,39)}$ = 19.3, p < 0.0001** | **$F_{(1,39)}$ = 21.5, p < 0.0001** |
| Day 5 Burying | 8.6 ± 3.0 | 111.5 ± 14.6$^{*}$ | 5.2 ± 1.4 | 38.8 ± 8.0$^{+}$ | **$F_{(1,40)}$ = 59.7, p < 0.0001** | **$F_{(1,40)}$ = 18.5, p = 0.0001** | **$F_{(1,40)}$ = 15.4, p = 0.0003** |
| Day 1 Rearing | 196.1 ± 33.5 | 261.8 ± 18.2 | 219.9 ± 35.7 | 235.3 ± 22.8 | $F_{(1,46)}$ = 2.0, p = 0.2 | $F_{(1,40)}$ = 0.002, p = 1.0 | $F_{(1,40)}$ = 0.8, p = 0.40 |
| Day 5 Rearing | 172.1 ± 15.7 | 225.4 ± 17.4$^{*}$ | 103.3 ± 22.5 | 228.4 ± 16.4$^{*}$ | **$F_{(1,42)}$ = 22.4, p < 0.0001** | $F_{(1,42)}$ = 5.0, p = 0.09 | $F_{(1,42)}$ = 3.6, p =0.06 |
| Day 1 LF | 99.9 ± 1.8 | 89.3 ± 8.5 | 119.1 ± 17.2 | 126.7 ± 15.8 | $F_{(1,23)}$ = 0.01, p = 0.91 | **$F_{(1,23)}$ = 3.2, p = 0.04** | $F_{(1,23)}$ = 0.5 p = 0.48 |
| Day 1 HF | 74.1 ± 10.0 | 40.6 ± 5.9 | 83.0 ± 14.8 | 69.6 ± 5.9 | **$F_{(1,21)}$ = 5.2, p = 0.03** | $F_{(1,21)}$ = 3.4, p = 0.08 | $F_{(1,21)}$ = 0.3, p = 0.34 |
| Day 1 LF/HF | 147.0 ± 21.9 | 183.7 ± 18.1 | 103.4 ± 7.5 | 126.5 ± 8.3 | **$F_{(1,22)}$ = 4.8, p = 0.04** | $F_{(1,22)}$ = 0.5, p = 0.49 | $F_{(1,22)}$ = 0.2, p = 0.67 |
| Day 5 LF | 130.8 ± 18.4 | 83.5 ± 6.9$^{*}$ | 65.6 ± 102.0 | 180.4 ± 125.0$^{+}$ | $F_{(1,23)}$ = 1.4, p = 0.25 | $F_{(1,23)}$ = 0.6, p = 0.46 | **$F_{(1,23)}$ = 11.7, p = 0.002** |
| Day 5 HF | 128.0 ± 40.2 | 30.2 ± 5.1$^{*}$ | 73.7 ± 15.4 | 75.3 ± 7.8 | **$F_{(1,22)}$ = 7.5, p =0.01** | $F_{(1,22)}$ = 0.1, p =0.80 | **$F_{(1,22)}$ = 8.0, p = 0.01** |
| Day 5 LF/HF | 130.7 ± 24.0 | 196.5 ± 20.28 | 123.5 ± 12.7 | 168.6 ± 20.1 | **$F_{(1,23)}$ = 6.4, p = 0.02** | $F_{(1,23)}$ = 0.6, p = 0.43 | $F_{(1,23)}$ = 0.2, p =0.64 |
| Sucrose Preference | 90.0 ± 2.8 | 77.9 ± 5.8 | 92.7 ± 1.7 | 82.3 ± 2.4 | **$F_{(1,43)}$ = 8.1, p = 0.007** | $F_{(1,43)}$ = 0.8, p = 0.37 | $F_{(1,43)}$ = 0.04, p = 0.83 |
| Plasma CORT | 85.8 ± 29.9 | 195.5 ± 35.5 | 146.6 ± 25.1 | 259.8 ± 56.0 | **$F_{(1,16)}$ = 7.7, p = 0.01** | $F_{(1,16)}$ = 2.3, p = 0.14 | $F_{(1,16)}$ = 0.002, p = 0.97 |

**Table S5. Summary of Statistical Statements for Study D.** Statistical analyses for effects of intra-CeA m-CLD are summarized in the table below. All data below were analyzed via 2-way ANOVA. Significant main effects are indicated in **bold** while significant post hoc analyses are denoted by the following: $^{*}p < 0.05$ vs drug-matched control; $^{+}p < 0.05$ vs stress condition-matched control

| Outcome of Interest | Vehicle Treatment | | Clodronate Treatment | | Statistical Analyses | | |
|---|---|---|---|---|---|---|---|
| | Control | Witness | Control | Witness | Stress | Treatment | Stress x Treatment Interaction |
| Day 1 Burying | 7.1 ± 1.5 | 65.9 ± 12.5* | 8.9 ± 2.9 | 100.4 ± 16.1* | **$F_{(1,47)} = 53.9, p < 0.0001$** | $F_{(1,47)} = 3.2, p = 0.08$ | $F_{(1,47)} = 2.5, p = 0.12$ |
| Day 5 Burying | 3.8 ± 1.1 | 86.1 ± 10.5* | 3.7 ± 1.0 | 98.8 ± 14.9* | **$F_{(1,48)} = 88.2, p < 0.0001$** | $F_{(1,48)} = 0.5, p = 0.51$ | $F_{(1,48)} = 0.5, p = 0.50$ |
| Day 1 Rearing | 288.4 ± 38.0 | 265.2 ± 17.0 | 276.8 ± 31.2 | 277.7 ± 16.0 | $F_{(1,49)} = 0.2, p = 0.68$ | $F_{(1,49)} = 0.0003, p = 0.99$ | $F_{(1,49)} = 0.2, p = 0.60$ |
| Day 5 Rearing | 237.4 ± 32.6 | 248.9 ± 23.3 | 263.3 ± 24.3 | 247.1 ± 24.5 | $F_{(1,50)} = 0.008, p = 0.90$ | $F_{(1,50)} = 0.2, p = 0.70$ | $F_{(1,50)} = 0.3, p = 0.60$ |
